# Multi-centre, multi-vendor reproducibility of 7T QSM and R_2_* in the human brain: results from the UK7T study

**DOI:** 10.1101/2020.04.22.055624

**Authors:** Catarina Rua, William T Clarke, Ian D Driver, Olivier Mougin, Andrew T. Morgan, Stuart Clare, Susan Francis, Keith Muir, Richard Wise, Adrian Carpenter, Guy Williams, James B Rowe, Richard Bowtell, Christopher T Rodgers

**Affiliations:** Wolfson Brain Imaging Centre, Department of Clinical Neurosciences, University of Cambridge, Cambridge, United Kingdom., (The Wolfson Brain Imaging Centre, Box 65, Cambridge Biomedical Campus, Cambridge, UK, CB2 0QQ); Wellcome Centre for Integrative Neuroimaging, FMRIB, Nuffield Department of Clinical Neurosciences, University of Oxford, Oxford, United Kingdom., (Wellcome Centre for Integrative Neuroimaging, FMRIB, Level 0, John Radcliffe Hospital, Oxford, United Kingdom, OX3 9DU); Cardiff University Brain Research Imaging Centre, School of Psychology, Cardiff University, Cardiff, United Kingdom., (Cardiff University Brain Research Imaging Centre, Cardiff University, Maindy Road, Cardiff, CF24 4HQ); Sir Peter Mansfield Imaging Centre, School of Physics and Astronomy, University of Nottingham, Nottingham, United Kingdom., (Sir Peter Mansfield Imaging Centre, University of Nottingham, University Park, Nottingham, NG7 2RD); Imaging Centre of Excellence, University of Glasgow, Glasgow, United Kingdom, (Imaging Centre of Excellence, Queen Elizabeth University Hospital, Langlands Dr, Glasgow, United Kingdom, G51 4LB); Department of Clinical Neurosciences and Cambridge University Hospitals NHS Trust, University of Cambridge, Cambridge, United Kingdom, (Department of Clinical Neurosciences, Herchel Smith Building, Cambridge Biomedical Campus, Cambridge CB2 0SZ); Medical Research Council Cognition and Brain Sciences Unit, University of Cambridge, Cambridge, United Kingdom., (MRC Cognition and Brain Sciences Unit, University of Cambridge, 15 Chaucer Road, Cambridge, CB27EF)

**Keywords:** 7 tesla, MRI, Quantitative Susceptibility Mapping, R_2_* mapping, Multi-centre, Reproducibility

## Abstract

We present the reliability of ultra-high field T_2_* MRI at 7T, as part of the UK7T Network’s “Travelling Heads” study. T_2_*-weighted MRI images can be processed to produce quantitative susceptibility maps (QSM) and R_2_* maps. These reflect iron and myelin concentrations, which are altered in many pathophysiological processes. The relaxation parameters of human brain tissue are such that R_2_* mapping and QSM show particularly strong gains in contrast-to-noise ratio at ultra-high field (7T) vs clinical field strengths (1.5 - 3T). We aimed to determine the inter-subject and inter-site reproducibility of QSM and R_2_* mapping at 7T, in readiness for future multi-site clinical studies.

**Methods:** Ten healthy volunteers were scanned with harmonised single- and multi-echo T_2_*-weighted gradient echo pulse sequences. Participants were scanned five times at each “home” site and once at each of four other sites. The five sites had 1x Philips, 2x Siemens Magnetom, and 2x Siemens Terra scanners. QSM and R_2_* maps were computed with the Multi-Scale Dipole Inversion (MSDI) algorithm (https://github.com/fil-physics/Publication-Code). Results were assessed in relevant subcortical and cortical regions of interest (ROIs) defined manually or by the MNI152 standard space.

**Results and Discussion:** Mean susceptibility (χ) and R_2_* values agreed broadly with literature values in all ROIs. The inter-site within-subject standard deviation was 0.001 – 0.005 ppm (χ) and 0.0005 – 0.001 ms^-1^ (R_2_*). For χ this is 2.1-4.8 fold better than 3T reports, and 1.1-3.4 fold better for R_2_*. The median ICC from within- and cross-site R_2_* data was 0.98 and 0.91, respectively. Multi-echo QSM had greater variability vs single-echo QSM especially in areas with large B_0_ inhomogeneity such as the inferior frontal cortex. Across sites, R_2_* values were more consistent than QSM in subcortical structures due to differences in B_0_-shimming. On a between-subject level, our measured χ and R_2_* cross-site variance is comparable to within-site variance in the literature, suggesting that it is reasonable to pool data across sites using our harmonised protocol.

**Conclusion:** The harmonized UK7T protocol and pipeline delivers on average a 3-fold improvement in the coefficient of reproducibility for QSM and R_2_* at 7T compared to previous reports of multi-site reproducibility at 3T. These protocols are ready for use in multi-site clinical studies at 7T.

## 1. Introduction

Neurodegenerative diseases are a significant global health burden. In many instances, neurodegeneration is associated with the deposition of iron in the brain. Understanding the patterns of deposition and their association with other risk factors is a key area of clinical research, but progress has been limited by the need to scale over multi-centre trials (Moeller et al., 2019). The EUFIND (Düzel et al., 2019) is an example of a network focused on advancements in neurodegenerative research by running large-scale multi-centre imaging studies. Also, the UK7T network (http://www.uk7t.org) has recently run a multi-site study with a dementia cohort to assess feasibility in patient groups. Imaging as part of the C-MORE study (Capturing the MultiORgan Effects of COVID-19) is also including harmonized multi-centre sequences which might provide insights into the long-term impact in survivors of COVID-19. Yet, in order to perform such multi-centre studies, it is necessary to first guarantee the consistency and reproducibility of imaging markers.

A popular approach to estimating iron concentration in the human brain uses gradient-echo (GE) magnetic resonance imaging (MRI). In grey matter, iron is mainly found in the protein ferritin which, due to its antiferromagnetic core and the presence of uncompensated spins at the surface or in the core, exhibits a superparamagnetic behaviour (Makhlof et al., 1997; Langkammer et al., 2012). This paramagnetic iron interacts with the MRI scanner’s static magnetic field (B_0_) causing local dipolar field perturbations. These accentuate the rate of transverse signal decay causing T_2_* relaxation in surrounding tissue, which is visible as decreasing signal amplitude with increasing echo time in a series of GE images. This effect causes an increase in the rate of transverse relaxation, R_2_*, which correlates well with non-heme iron concentrations in grey matter (Gelman et al., 1999; Langkammer et al., 2010), and has been used to investigate the distribution of iron in the healthy brain and in disease (Haacke et al., 2005; Yao et al., 2009; Li et al., 2019).

The local presence of iron (and to a lesser extent myelin and calcium) also affects the signal phase of GE images because of the effect of the field perturbation on the local Larmor frequency (House et al., 2007; He et al., 2009; Lee et al., 2012). Quantitative Susceptibility Mapping (QSM) methods attempt to deconvolve these dipole phase patterns to identify the sources of the magnetic field inhomogeneity. In other words, QSM estimates quantitative maps of tissue magnetic susceptibility χ from GE phase data (Li and Leigh, 2004; Reichenbach, 2012; Wang and Liu, 2015). This approach has shown sensitivity to several neurological conditions (Lotfipour et al., 2012; Acosta-Cabronero et al., 2013; Blazejewska et al., 2015; Acosta-Cabronero et al., 2016) and offers advantages over magnitude R_2_* such as having reduced blooming artifacts or being able to distinguish between paramagnetic and diamagnetic substances (Eskreis-Winkler et al., 2017).

R_2_* imaging and QSM have been shown to provide reproducible results in single-site and cross-site studies at 1.5T and 3T (Hinoda et al., 2015; Cobzas et al., 2015; Deh et al., 2015; Lin et al., 2015; Santin et al., 2017; Feng et al., 2018; Spincemaille et al., 2019).

The dipole-inversion problem at the heart of QSM methods benefits from the increased sensitivity to magnetic susceptibility variation and spatial resolution at ultra-high fields (B_0_ ≥ 7 T) (Yacoub et al., 2001; Reichenbach et al., 2001; Tie-Qiang et al., 2006; Duyn et al., 2007; Wharton and Bowtel, 2010). At 7T, close attention must be paid to B_0_ shimming and gradient linearity to achieve accurate QSM and R_2_* mapping (Yang et al., 2010). Head position is also an important factor that affects the susceptibility anisotropy (Lancione et al., 2017; Li et al., 2017).

In this study, we introduce single-echo and multi-echo GE imaging protocols for QSM and R_2_* mapping at 7T which were standardised on three different 7T MRI scanner platforms, from two different vendors. We applied this standardised protocol in the UK7T Network’s “Travelling Heads” study on 10 subjects scanned at 5 sites. We report reproducibility for derived R_2_* and QSM maps and make recommendations for the design of future multi-centre studies.

## 2. Methods

### 2.1. Measurement setup

Ten healthy volunteers (3 female, 7 male; age 32.0±5.9 years) were recruited: comprising two subjects from each of the five 7T imaging sites in the UK7T Network (described in Table 1). Each subject was scanned five times at their “home” site, and once at the other sites, under local ethics approval for multi-site studies obtained at Site-4 (HBREC.2017.08). Scans for each subject were completed within a period of between 83 and 258 days. The five home-site scans were performed across different sessions: the median time to acquire all five scans was 59 days (range: 3-71 days).

**Table 1:** Details of the scanners and hardware used for the UK7T Network’s Travelling Heads study.

| # Site | Vendor | Scanner Model | Gradient Performance | Installation Date (Month-Year) | Software Version |
| --- | --- | --- | --- | --- | --- |
| 1 Wellcome Centre for Integrative Neuroimaging (FMRIB), University of Oxford | Siemens | Magnetom 7T | 70 mT m<sup>-1</sup><br>200 mT m<sup>-1</sup> ms<sup>-1</sup> | Dec-2011 | VB17A |
| 2 Cardiff University Brain Research Imaging Centre, Cardiff University | Siemens | Magnetom 7T | 70 mT m<sup>-1</sup><br>200 mT m<sup>-1</sup> ms<sup>-1</sup> | Dec-2015 | VB17A |
| 3 Sir Peter Mansfield Imaging Centre, University of Nottingham | Philips | Achieva 7T | 40 mT m<sup>-1</sup><br>200 mT m<sup>-1</sup> ms<sup>-1</sup> | Sep-2005 | R5.1.7.0 |
| 4 Wolfson Brain Imaging Centre, University of Cambridge | Siemens | Magnetom Terra | 80 mT m<sup>-1</sup><br>200 mT m<sup>-1</sup> ms<sup>-1</sup> | Dec-2016 | VE11U |
| 5 Imaging Centre of Excellence, University of Glasgow | Siemens | Magnetom Terra | 80 mT m<sup>-1</sup><br>200 mT m<sup>-1</sup> ms<sup>-1</sup> | Mar-2017 | VE11U |

In every scan session, B_0_ shimming was performed using the vendors’ default second-order (or third-order for Site-4 and Site-5) B_0_-shimming routines. B_1_^+^-calibration was performed initially using the vendor’s default adjustment scans. A 3D DREAM sequence (Nehrke et al., 2012; Ehses et al., 2019) was subsequently acquired and the transmit voltage (or power attenuation) was then adjusted for all subsequent imaging based on the mean flip-angle from the brain in an anatomically-specified axial slice of the 3D DREAM flip angle map as described in Clarke et al. (2019). Single-echo 0.7mm isotropic resolution T_2_*-weighted GE data were then acquired with: TE/TR=20/31ms; FA=15°; bandwidth=70Hz/px; in-plane acceleration-factor=4 (Sites-1/2/4/5) or 2×2 (Site-3); FOV=224×224×157mm^3^; scan-time=∼9min. Multi-echo 1.4mm isotropic resolution T_2_*-weighted GE data were acquired with: TE_1_/TR=4/43ms; 8 echoes with monopolar gradient readouts; echo-spacing=5ms; FA=15°; bandwidth=260Hz/px; acceleration-factor=4 (Sites-1/2/4/5) or 2×1.5 (Site-3); FOV=269×218×157mm^3^; scan-time ∼6min (Sites-1/2/4/5) and ∼4min (Site-3). For Siemens data, coil combination was performed using a custom implementation of Roemer’s algorithm, as previously described (Clarke et al., 2019). Subject 6’s single-echo scan failed to reconstruct using Roemer’s method on data from the 1^st^ visit at Site-5 so a sum-of-squares (SoS) algorithm was used for coil combination for that scan instead. A 0.7mm isotropic MP2RAGE scan was used for within-and cross-site registration as previously described (Mougin et al., 2019).

### 2.2. QSM and R_2_* data processing

QSM maps were generated from both the single-echo and multi-echo T_2_*-weighted datasets using the Multi-Scale Dipole Inversion (MSDI) algorithm, as implemented in QSMbox v2.0 (Acosta-Cabronero et al., 2018). Briefly: first the local field was estimated by phase unwrapping (Abdul-Rahman et al., 2005) and magnitude-weighted least squares phase echo fitting was performed on the multi-echo data. Then, independently for both single-echo and multi-echo data, background field was removed using the Laplacian Boundary Value (LBV) method followed by the variable Spherical Mean Value (vSMV) algorithm with an initial kernel radius of 40mm (Zhou et al., 2014; Acosta-Cabronero et al., 2018). MSDI inversion was estimated with two scales: the self-optimised lambda method was used on the first scale with filtering performed using a kernel with 1mm radius, and on the second scale the regularization term was set to λ=10^2.7^ (the optimal value for *in-vivo* 7T datasets found in (Acosta-Cabronero et al., 2018)) and filtering was done with a kernel radius set to 5mm. Brain masks used in the analysis were obtained with FSL’s Brain Extraction Tool (BET) with fractional intensity threshold=0.2 for single-echo data (Smith, 2002). These were then mapped to multi-echo data space.

On the multi-echo data, QSM was reconstructed seven more times: with only one echo at 19 ms (matching the single echo data), with the two shortest echoes (i.e. TE_1_/TE_2_ = 4/9 ms), with the three shortest echoes (i.e. TE_1_/TE_2_/TE_3_ = 4/9/14 ms), and so forth. On the multi-echo dataset, voxel-wise quantitative maps of R_2_* were obtained using the Auto-Regression on Linear Operations (ARLO) algorithm for fast monoexponential fitting (Pei et al., 2015). R_2_* was also fitted five more times: with data from the first three echoes (TE1/TE2/TE3=4/9/14 ms), then with the first four echoes (TE1/TE2/TE3/TE4=4/9/14/19 ms), and so forth.

### 2.3. Data Registration

The neck was cropped from the magnitude data with FSL’s “robustfov” command (https://fsl.fmrib.ox.ac.uk/fsl/), applied to the single-echo data and the 4^th^ echo of the multi-echo data. High-resolution single-echo and multi-echo templates were made fromthiscroppeddataforeachsubjectwith antsMultivariateTemplateConstruction2.sh from the Advanced Normalization Tools (ANTs, http://stnava.github.io/ANTs/). Two approaches were compared: transformations using rigid registration with mutual information similarity metric (denoted as “Rigid” below) or using symmetric diffeomorphic image registration with cross-correlation similarity metric (denoted “SyN” below). Other settings were kept the same for both approaches: 4 steps with 0.1 gradient step size, maximum iterations per step 1000, 500, 250 and 100, smoothing factors per step of 4, 3, 2, and 1 voxels, and shrink factors per step of 12x, 8x, 4x, and 2x. The resulting registrations were then applied to the QSM and R_2_* maps which were averaged to create single-echo and multi-echo QSM and R_2_* templates for each subject.

### 2.4. Selection of Regions of Interest (ROIs)

Five regions of interest (Substantia Nigra, Red Nucleus, Caudate Nucleus, Putamen and Globus Pallidus) were manually segmented based on the subject-specific QSM templates of the single-echo data registered with the “SyN” approach. In order to minimize the amount of segmentation variability, these ROIs were then mapped to the single-echo “Rigid”, and multi-echo “SyN” and multi-echo “Rigid” spaces with nearest neighbour interpolation and via non-linear registrations obtained with the default settings in the antsRegistrationSyN.sh command in ANTs.

Magnitude data were first registered to the T_1_-weighted MP2RAGE scans (Rigid transformations; MI similarity metric) and later to the standard T_1_ “MNI152 brain” (Montreal Neurological Institute 152) (using settings in antsRegistrationSyN.sh) applied to the single-echo data and to the 1^st^ echo of the multi-echo data. These registrations were then used to map the 48 probabilistic cortical ROIs, “cortical ROIs”, from the Harvard-Oxford Cortical Atlas and the 21 probabilistic subcortical ROIs, “subcortical ROIs”, from the Harvard Oxford Subcortical Atlas to the QSM and R_2_* template spaces. The T_1_-weighted MP2RAGE data was bias-field corrected, brain extracted, and segmented into five tissues using SPM (https://www.fil.ion.ucl.ac.uk/spm/): the grey matter (GM), white matter (WM) and cerebral-spinal fluid (CSF) volumes were mapped into each subject-specific QSM template space. Then, using “fslmaths” from FSL (https://fsl.fmrib.ox.ac.uk/fsl/), the mapped cortical ROIs were thresholded at 10% of the “robust range” of non-zero voxels and multiplied by the GM tissue map in order to obtain GM-specific cortical ROIs. The mapped subcortical ROIs were thresholded at 50% of the “robust range” of non-zero voxels. From these, any CSF voxels were excluded from the left and right Caudate Nucleus, Putamen and Globus Pallidus, and the voxel sets from the left and right counterparts were merged together.

From the single-echo and multi-echo data, average χ and R_2_* values were extracted from the manual and Atlas-based ROIs for all volunteers and sessions in template space (values given in Supplementary Material 1).

were first estimated from the multi-echo datasets. Δ *B*_0_was calculated from the In order to estimate where the magnetic field is spatially more variable, field-maps background field removal step of the QSM pipeline, and was defined, per-voxel, as the The average Δ *B*_0_ was extracted for each of the cortical ROIs and averaged across all average difference between the field in a voxel and its immediate nearest neighbors. the Δ *B*_0_ values: wherever | Δ *B*_0_|>0.005 *HZ* the ROI was grouped into “high Δ *B*_0_ “subjects and sessions. Then the cortical ROIs were divided into two groups based on regions, otherwise it was grouped into “low Δ *B*_0_ “regions. Δ *B*_0_” was calculated from the multi-echo pipeline only, as differences to values calculated using single-echo data were minimal (Figure 1, Supplementary Material 2).

**Figure 1:**
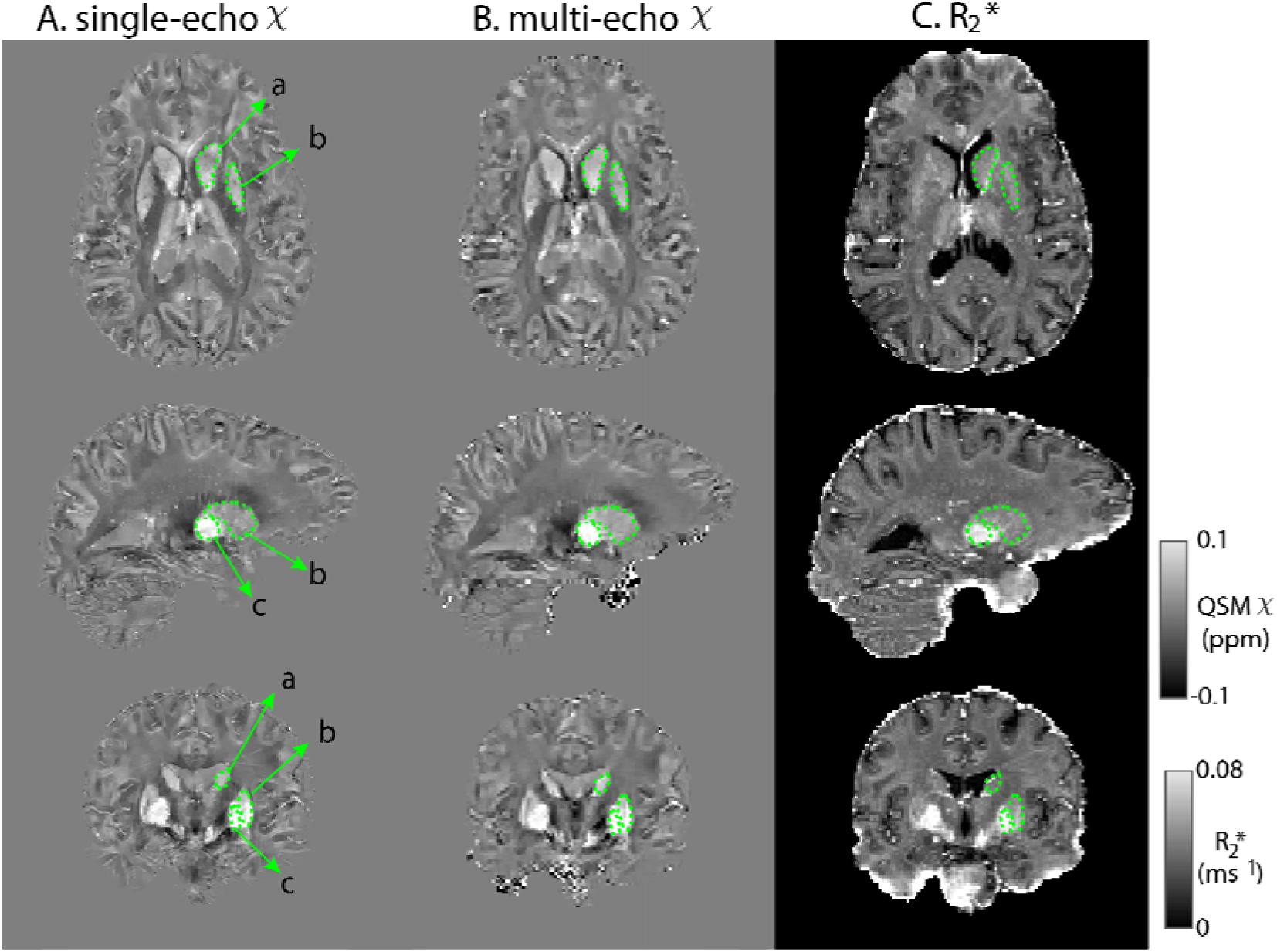
Representative slices of single-echo χ (A) multi-echo χ (B) and R_2_* maps (C) from an example subject templates. The right Caudate Nucleus (a), Putamen (b) and Globus Pallidus (c) are shown in green. Multi-echo χ maps calculated with data from all 8 echoes.

We explored three possible susceptibility reference regions for QSM processing. The average QSM signal was extracted from:

1. A whole brain mask, “wb”;
2. A whole-brain CSF mask eroded in two steps, “csf”;
3. A manually placed cylindrical ROI in the right ventricle, “cyl” (across all subjects the ROI volume was 104±11 mm^3^).

### 2.5. Statistical Analysis

Statistical analysis was performed with R 3.5.3 (R Core Team, 2013). Cross-site analysis used only the 1^st^ scan at the “home” site along with the scans at the other four sites. To obtain the within subject average, AV_w_, the χ and R_2_* values were averaged within the same site and across the sites and then averaged across subjects:

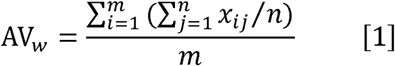

where *n* is the number of sessions (*n* = 5 hin-site and cross-site) and *m* the number of subjects. Relative reliability was measured using the intra-class correlation coefficient (ICC) from within and cross-site data independently for each ROI (Weir, 2005):

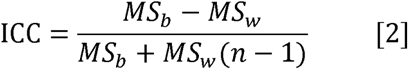

where *MS*_*b*_ and *MS*_*w*_ are the between-subjects and within-subjects mean square from a random-effects, one-way analysis of variance (ANOVA) model. Intra-subject absolute variability is assessed by measuring the within-subject standard-deviation (SD_w_) calculated as (Santin et al., 2017):

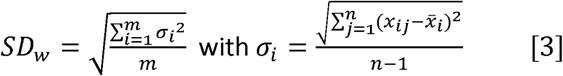

where 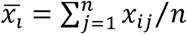 is the replicate average for each subject. SD_w_ was computed using within-site data and cross-site data independently. Similarly, cross-subject variability was calculated by measuring the between-subject standard-deviation (SD_b_):

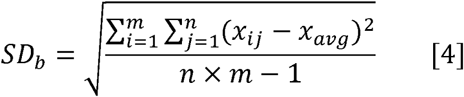

where 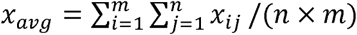 is the measurement average across subjects and sessions. Note that SD_b_ is computed using data from all sites.

Statistical testing on AV_w_, SD_w_ and ICC values extracted from manual and template-based ROIs was done by first fitting the data with normal, log-normal, gamma and logistic distributions. The goodness-of-fit statistics for the parametric distributions were calculated and the distribution which showed the lowest Akaikes Information Criterion (AIC) was then used on a general linear model fitting. All models included as fixed main effects ROI number and data type (within-and cross-site). When evaluating the data registration type, the model also included registration type (“Rigid” and “SyN”) as a fixed main effect. When testing for QSM reference, the model also included reference region (“wb”, “csf”, and “cyl”) as a fixed main effect. On multi-echo QSM data, a model was fitted which also included the number of echoes processed as a fixed main effect. When comparing the manual and subcortical ROIs, the ROI type from the cortical ROIs, ROI number was replaced with “high Δ *B*_0_ “and “low Δ *B*_0_ “ROI (manual vs. atlas-based) was also included as a fixed main effect. Finally, on the data type as covariate. A p-value less than 0.05 was considered significant.

### 2.6. Head orientation

We investigated the effect of head orientation on QSM variability. Since all our data was acquired with transverse slice orientation, the slice normal vector in the acquired images was parallel to B_0_. We used the per-subject rotation matrices of the affine transforms from this acquired transverse data to MNI space to estimate the z-axis rotation θ with respect to the B_0_ vector (0,0,1) (Figure 7 (A)):

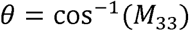

where *M*_33_ is the 3^rd^ row, 3^rd^ column of the affine transform matrix.

Two linear mixed effects models, ‘mod1’ and ‘mod2’, were fitted on the within-site and cross-site χ data separately: both models included site, ROI, and session number Two linear mixed effects models, ‘mod1’ and ‘mod2’, were fitted on the within-site as fixed effects, and subject number as a random effect, while ‘mod2’ also included *θ* as a fixed effect. For each model, the R^2^ was evaluated and both models were compared with a chi-squared test.

Finally, from ‘mod2’ the *θ* fit coefficients were used to estimate corrected χ -values based on a chosen standard *θ* for all of the measurements (*θ* _*norm*_ *=* 0.52 radians).

Then, new within-site and cross-site SD_w_ of the corrected were calculated based on the same approach as in sub-section 2.5.

## 3. Results

Figure 1 shows QSM and R_2_* maps for one example subject. Basal ganglia structures, including Caudate Nucleus, Putamen and Globus Pallidus are clearly visible consistent with previous findings (Langkammer et al., 2010; Wang et al., 2015; Betts et al., 2016; Acosta-Cabronero et al., 2016). Figure 2, Supplementary Material 2 highlights the difference in QSM data quality when using our chosen Roemer coil combination method vs using sum-of-squares coil combination.

**Figure 2:**
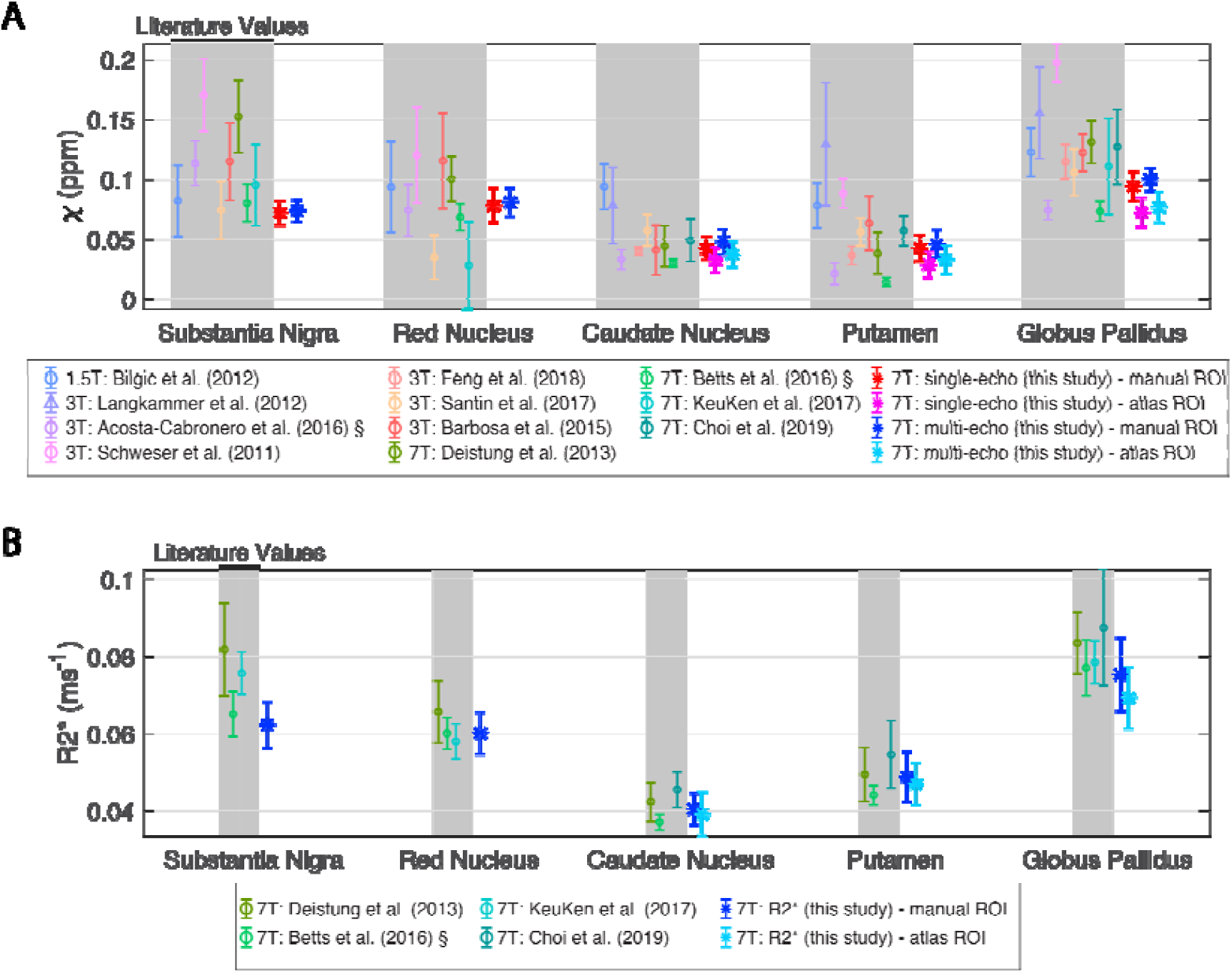
Mean and standard deviation literature values of QSM (A) and R_2_* (B). The mean and standard deviation results from this study are also plotted. For data with the symbol ‘§’ the standard error of the mean was originally reported and has been rescaled by reported N. Shaded regions correspond to literature data. Multi-echo χ-maps were calculated with data from all eight echoes.

### 3.1. QSM and R_2_* results and literature

Figure 2 compares average χ and R_2_* values calculated in this study in the five manual ROIs and three corresponding atlas-based subcortical ROIs against literature ranges. The single-echo χ-values and multi-echo χ-values from this study are consistent with literature values at 1.5T, 3T and 7T. R_2_* values from this study also agree closely with 7T literature values.

### 3.2. Reproducibility of QSM and R_2_*

Figure 3 shows boxplots over ROIs of the within- and cross-site AV_w_ (A), SD_w_ (B) and ICC (C) values for the manual ROIs on the χ and R_2_* maps. The AV_w_ from R_2_* maps measured on the same site is systematically higher compared to the AV_w_ measured across sites (p < 0.0001; e.g., on the Putamen ROI, AV_w_within-site_ = 0.0493 ms^-1^ vs AV _w_cross-site_ = 0.0489 ms^-1^). On this comparison, QSM data did not show significant differences between within-site and cross-site groups for either single-echo data (p = 0.053) or multi-echo data (p = 0.65).

**Figure 3.**
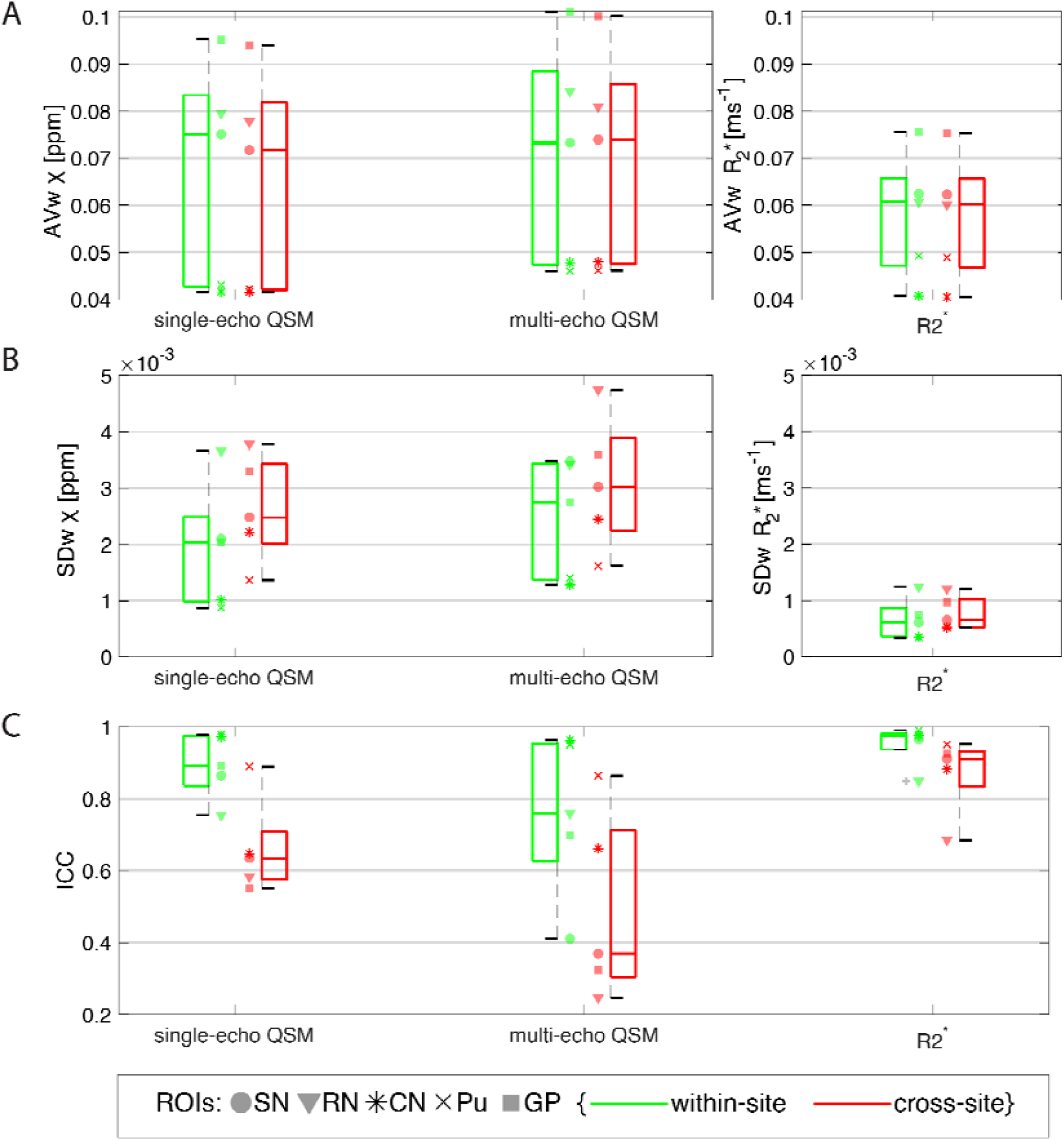
Boxplots from data obtained on the manual ROIs of within- and cross-site AV_w_ (A), SD_w_ (B) and ICC (C) of single-echo and multi-echo QSM, and R_2_*. Data from each ROI is shown with a different marker for each boxplot. Legend: SN=Substantia Nigra; RN: Red Nucleus; CN: Caudate Nucleus; Pu: Putamen; GP: Globus Pallidus. The variability in AV_w_ reflects the natural variation of iron content in subcortical structures in the healthy brain. Multi-echo χ-maps were calculated with data from all eight echoes.

From all the data in the manual ROIs, the median SD_w_ of single-echo χ-values was approximately 29% lower than for multi-echo χ-values (p = 0.0010). There was a significantly larger SD_w_ cross-site compared to within-site on single-echo χ data (p < 0.0001; e.g., on the PN ROI, SD_w_within-site_ = 0.00088 ppm vs SD_w_cross-site_ = 0.0014 ppm), multi-echo χ (p = 0.033) and on R_2_* data (p < 0.0001).

The ICC values for within- and cross-site R_2_* data (median ICC was 0.98 and 0.91, respectively) were found to be significantly higher than values for single-echo χ (median ICC was 0.89 and 0.64, respectively) or for multi-echo χ (median was ICC 0.76 and 0.38, respectively) (p = 0.00011). For all measurements, the ICC for cross-site data was significantly lower than for within-site data (single-echo QSM: p < 0.0001; multi-echo QSM: p = 0.017; R_2_*: p < 0.0001).

Similar statistics were obtained for AV_w_, SD_w_ and ICC measurements in the Altas-based cortical ROIs (Table 2, Supplementary Material 2).

**Table 2:** Within-site to Cross-site ratio of the median SD_w_ obtained from all five manually-defined ROIs on single-echo and multi-echo *χ* without and with *θ*-correction.

| Parameter | within-site/cross-site median $SD_w$ (no $\theta$ - correction) | within-site/cross-site median $SD_w$ (with $\theta$ - correction) |
| --- | --- | --- |
| Single-echo $\chi$ | 0.82 | 0.67 |
| Multi-echo $\chi$ | 0.91 | 0.82 |

### 3.3 Registration

The within- and cross-site standard deviations for one axial slice from one example subject using “Rigid” and “SyN” registration approaches are shown in Figure 4. Generally, with both registration methods, within-site and cross-site SD_w_ increases in veins, in the orbitofrontal regions and at the cortical surface (white and green arrows, Figure 4). These are areas associated with large B_0_ inhomogeneities and gradient non-linearity. However, there is a decrease in the cross-site standard deviation in the orbitofrontal region and close to the edges of the cortex when using the “SyN” compared to the “Rigid” method (green arrows, Figure 4).

**Figure 4.**
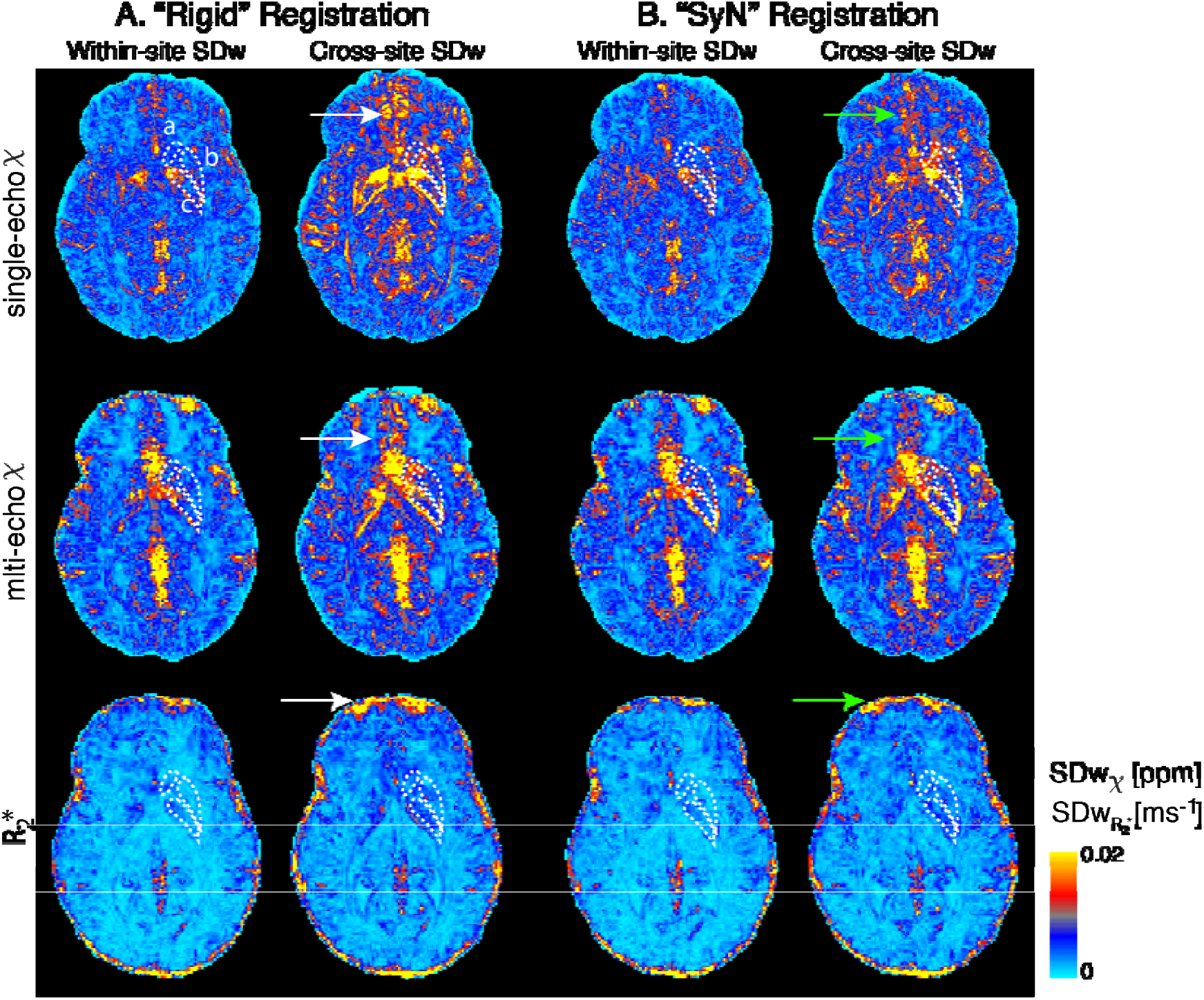
Voxel-wise within- and cross-site standard deviation of an example subject from single-echo and multi-echo QSM and R_2_* data with data registered with “Rigid” (A) and “SyN” (B) transformations. Arrows point to regions where the SD_w_ decreased with the “SyN” transformations (green) are compared to “Rigid” (white). The right Caudate Nucleus (a), Putamen (b) and Globus Pallidus (c) are outlined in white. Multi-echo χ-maps were calculated with data from all eight echoes.

On the manual ROIs increased variability was observed for R_2_* on “Rigid” registered data compared to “SyN” (SD_w_: p < 0.0001; ICC: p < 0.013) but not for single-echo or multi-echo χ: for example, the median cross-site R_2_* SD_w_ from all ROIs was 0.00066 ms^-1^ using “SyN” method and 0.00086 ms^-1^ using the “Rigid” registration method. On the atlas-based cortical ROIs, the same significant trend was observed for R_2_* and single-echo χ data (Table 2, Supplementary Material 2).

### 3.4 QSM referencing

To assess the optimal QSM susceptibility referencing, Figure 5 shows boxplots of the SD_w_ for single-echo and multi-echo χ using different referencing methods on the manual ROIs. On single-echo χ data, compared to “wb” correction (chosen correction for this study), the “csf” reference did not increase significantly the SD_w_ (p = 0.93) but with “cyl” the median SD_w_ increased by approximately 14% (p < 0.0001).

**Figure 5:**
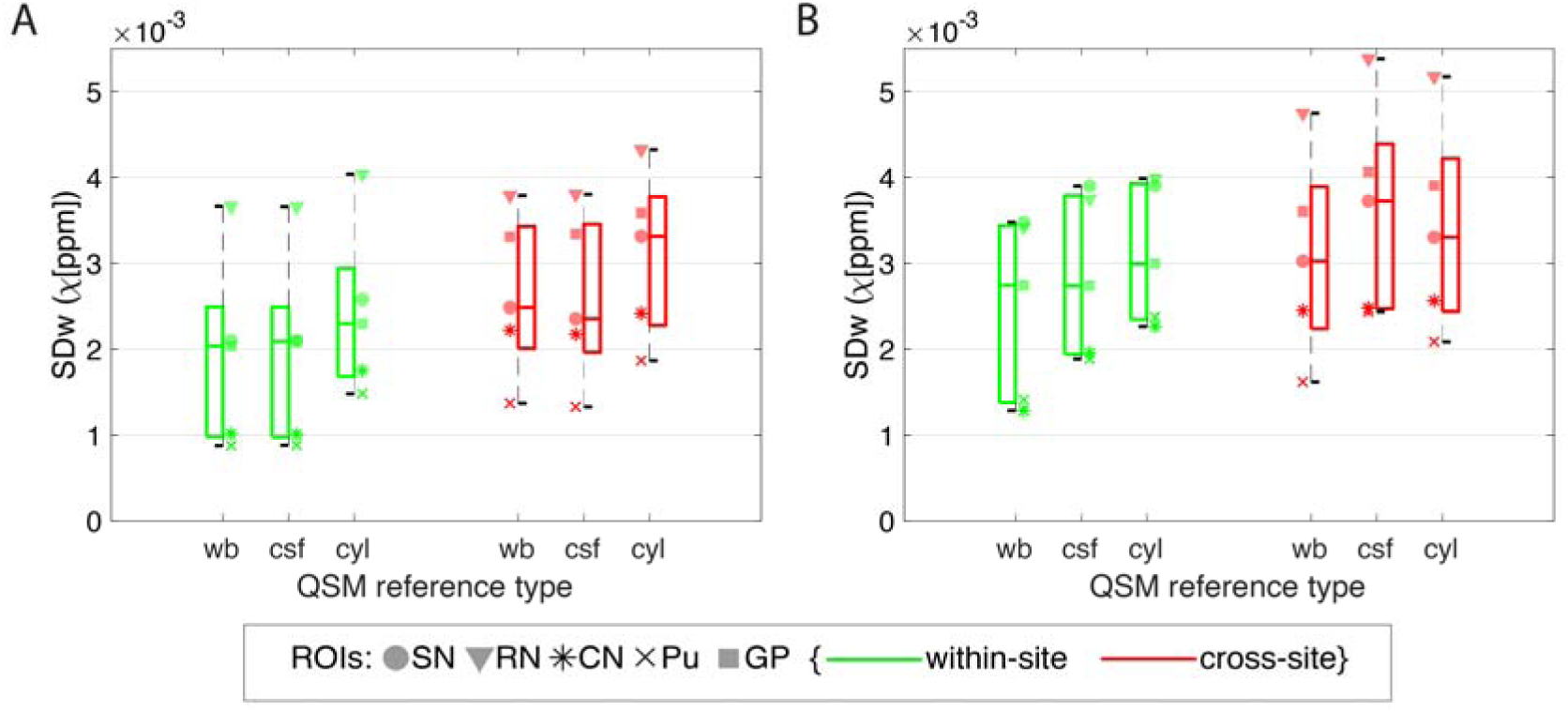
Boxplots from data obtained on the manual ROIs of within- and cross-site SD_w_ (red and green, respectively) of single-echo QSM (A) and multi-echo QSM (B) with a whole-brain reference (wb), with a csf reference (csf), and with a cylinder reference (cyl). Data from each ROI is shown with a different marker for each boxplot. Legend: SN=Substantia Nigra; RN: Red Nucleus; CN: Caudate Nucleus; Pu: Putamen; GP: Globus Pallidus. Multi-echo χ-maps were calculated with data from all eight echoes.

multi-echo χ data showed an increase in the median SD_w_ of, respectively, 11% (p = 0.00096) and 8% (p = 0.00064) when using “csf” and “cyl” methods for correction. The effect of varying the referencing of QSM data was similar in within-site and cross-site data, for all methods tested.

### 3.5 Multi-echo QSM & R2*

On average across all the manual ROIs and compared to single echo data, multi-echo data (using two or more echoes) showed a significant 14% increase of the SD_w_ (Figure 6) and 3% of the ICC (Table 1, Supplementary Material 2). This supports the single-echo and multi-echo χ comparison in Section 3.2. Similar behaviour was observed on the atlas-based cortical ROIs (Table 2, Supplementary Material 2). On the manual ROIs, there is no significant difference in AV_w_ (p = 0.79) or in SD_w_ (p = 0.11) from χ computed from multiple echoes (i.e. 2 or more echoes in the QSM analysis). Yet, in the atlas-based cortical ROIs, long echo times (i.e. using 6 or more echoes) showed an average increase of 15.7% in SD_w_ (p < 0.0001) compared to using 2 to 5 echoes and a decrease of 1.75% in ICC (p < 0.0001) (Table 2, Supplementary Material 2).

**Figure 6.**
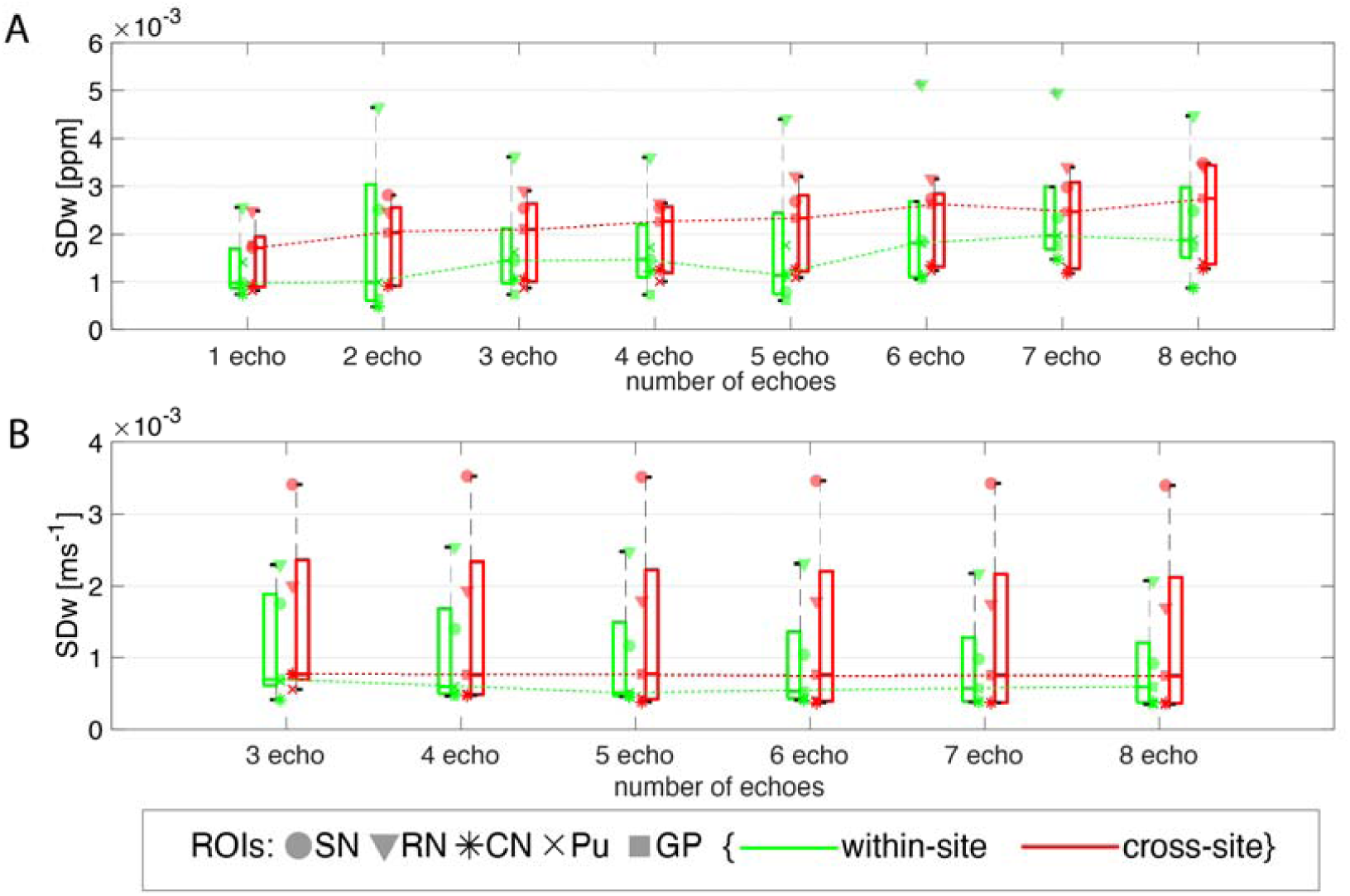
Boxplots from data obtained on the manual ROIs of within- and cross-site SD_w_ for multi-echo QSM (A) and R_2_* (B) calculated with different number of echoes. Increasing trend on the median SD_w_ observed with increasing number of echoes was observed on the QSM data (dotted green and red dotted lines in (A)). Legend: SN=Substantia Nigra; RN: Red Nucleus; CN: Caudate Nucleus; Pu: Putamen; GP: Globus Pallidus.

In the manual ROIs, R_2_* showed no significant change in variability across all ROIs when different number of echoes were used in the fitting (SD_w_: p = 0.11; ICC: p = 0.95) (Figure 6 (B)) or on AV_w_ (p = 0.97). In the atlas-based cortical ROIs, the number of echoes used influenced the average R_2_* value (AV_w_: p < 0.0001), weakly ICC (p = 0.021), but not SD_w_ (p = 0.61). Table 1 and 2, Supplementary Material 2 display individual statistics.

### 3.6 ROI selection

There is a small but significant higher average χ from manually drawn ROIs compared to the atlas-based subcortical ROIs in single-echo QSM data (p < 0.0001; e.g. 0.042±0.009 ppm vs 0.033±0.010 ppm in the caudate nucleus) and in multi-echo QSM data (p < 0.0001; e.g. 0.048±0.010 ppm vs 0.038±0.011 ppm in the caudate nucleus) (Figure 2). Similarly, for R_2_* (e.g. 0.041±0.004 ms^-1^ vs 0.039±0.006 ms^-1^ in the caudate nucleus) this difference was significant (p < 0.0001). In addition, the SD_w_ was, on average, approximately two times higher and the ICC lower in the atlas-based subcortical ROIs compared to the manual ROIs in all datasets (SD_w_: single-echo QSM p < 0.0001, multi-echo QSM p < 0.0001, R_2_* p < 0.0001; ICC: single-echo QSM p = 0.00021, multi-echo QSM p = 0.0023, R_2_* p = 0.012). So, ROI selection should be done consistently in a study.

### 3.7 Spatial distribution of the magnetic field

On the altas-based cortical ROIs the SD_w_ increased by approximately 28% and 88% on “high Δ*B*_0_” regions compared to “low Δ*B*_0_” regions on multi-echo χ and R_2_* data, respectively (p = 0.0011 and p < 0.0001) (Table 2, Supplementary Material 2). Similarly, ICC values decreased significantly for single-echo and multi-echo χ and R_2_* values.

### 3.8 QSM variability with head orientation

When analysing χ in the manually-defined ROIs with respect to *θ*, a consistent negative trend was observed for all subjects. Figure 7 (C and D) show an example for the analysis in the Globus Pallidus ROI. Fitting a linear model on χ χ, with *θ* and ROI as fixed variables, *θ* showed a significant negative correlation with single-echo χ (p < 0.0001) and multi-echo χ (p = 0.015).

**Figure 7.**
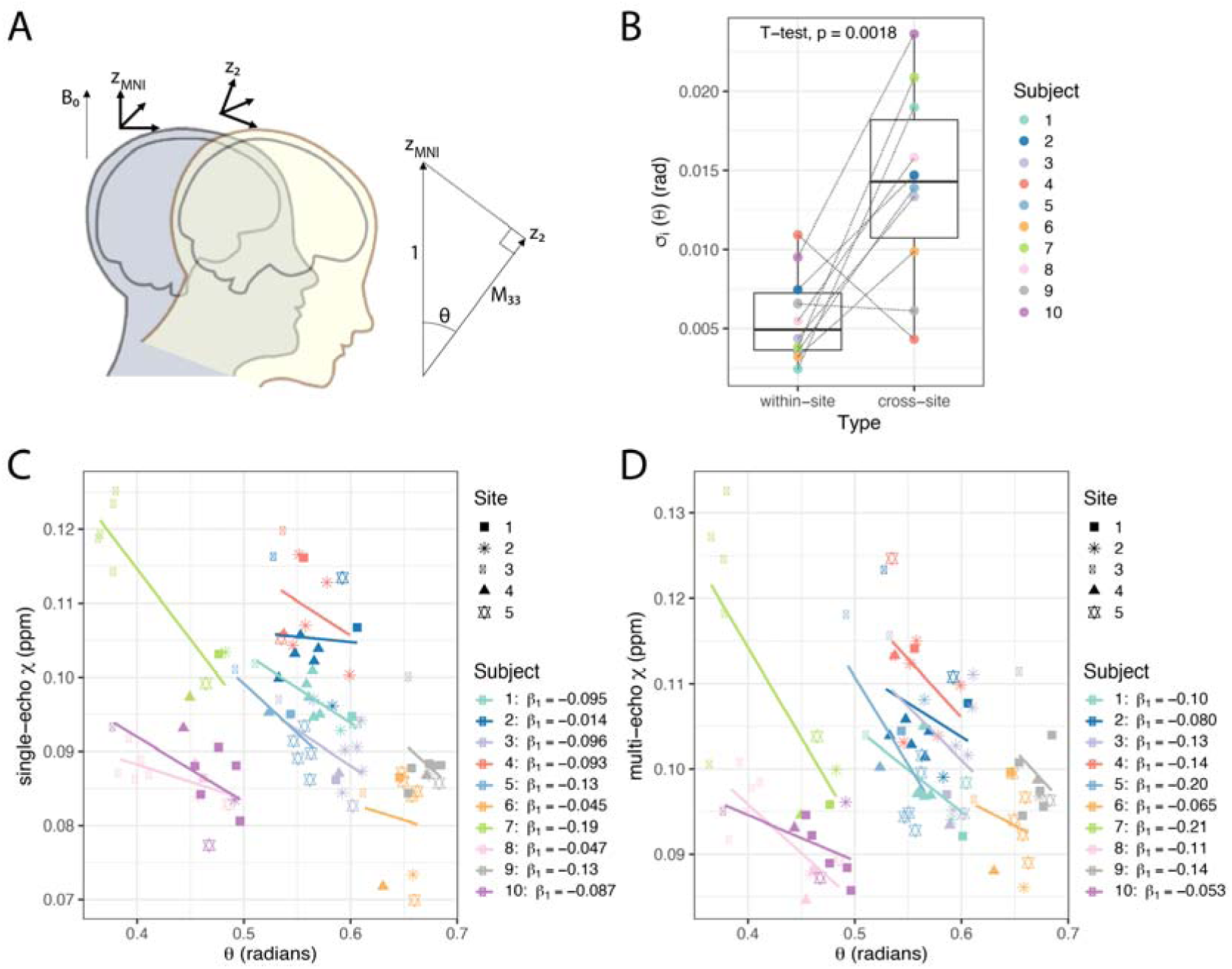
In QSM, it is assumed that the macroscopic susceptibility in an imaging voxel is isotropic. However, it has been shown that this assumption is too simplistic for single head orientation QSM methods, complicating the interpretation of the QSM results (Li et al., 2017). We investigated the effect of head orientation on QSM estimation in our data: (A) Considering that data was all acquired in the transverse plane with B_0_ perpendicular to the imaging slice, subjects had a variable head rotation *θ* with respect to B_0_. To estimate *θ*, we used MNI space as a common head orientation (z_MNI_) across all scans. From the affine registration matrix converting acquired data into MNI space, the angle of rotation from the rotated z-axis, z_2_, will be given by *θ*= cos^−1^ (*M*_33_) where *M*_33_ is the 3^rd^ row, 3^rd^ column of the affine transform matrix. (B) Subject-wise within-site and cross-site (J_l_ measurements on 0. (C) Single-echo and (D) multi-echo scatter plots of x measurements according to 0 on the Globus Pallidus manual ROI. For each subject a linear trend is also plotted and the fit coefficients are given in the plot legend. Data from each site is displayed with a different symbol. Multi-echo χ-maps were calculated with data from all eight echoes.

In addition, for *θ* the within-site SD_w_ was nearly half of the cross-site SD_w_ (0.011 and 0.028 radians, respectively), indicating that there was larger variability in head orientation across sites (subject-wise variability of *θ*, σ_*i*_ of equation [3], is plotted in Figure 7 (B)).

Separately for within-site and cross-site χ data, we assessed the goodness-of-fit of a model containing *θ* as an explanatory variable. On single-echo within-site data, the marginal R^2^ increased from 0.71 with ‘mod1’ to 0.76 with ‘mod2’ (which includes *θ*) (Chi-squared test, p = 0.041). The corresponding cross-site R^2^s were: 0.77 and 0.80 (Chi-squared test, p = 0.057). On multi-echo data, the marginal R^2^ increased from 0.75 with ‘mod1’ to 0.79 with ‘mod2’ on within-site data (Chi-squared test, p = 0.041) and maintained at 0.79 on both models for cross-site data (Chi-squared test, p = 0.14).

From the corrected x-values at *θ*_*norm*_, results show a slight decrease in the ratio of within-site to cross-site SD_w_ (Table 2), but variability of χ obtained from cross-site data was still higher than from within-site (x with *θ*-correction, p=0.01; uncorrected χ, p < 0.0001 (subsection 3.2)). For multi-echo data, the SD_w_ obtained from the corrected χ-values were similar on within-site compared to cross-site (χ with *θ*-correction, p=0.11; uncorrected χ, p = 0.033 (subsection 3.2)).

## 4. Discussion

In this paper, the reproducibility of QSM χ and R_2_* measurements in cortical and subcortical regions of the brain was assessed for the first time in a multi-site study at 7T for two different protocols (a single-echo 0.7mm isotropic T_2_*-weighted scan and a 1.5mm isotropic multi-echo T_2_*-weighted scan), using three different scanner platforms provided by two different vendors.

Previous studies at 1.5T and 3T have shown good reproducibility for χ and R_2_* data acquired on the same scanner or across sites (1.5T and 3T) (Hinoda et al., 2015; Cobzas et al., 2015; Deh et al., 2015; Lin et al., 2015; Santin et al., 2017; Feng et al., 2018; Spincemaille et al., 2019). In terms of QSM and depending on the subcortical region, intra-scanner 3T repeatability studies report an SD_w_ of 0.002-0.005 ppm (Feng et al., 2018) and 0.004-0.006 ppm (Santin et al., 2017), and the cross-site 3T study by Lin et al. (2015) reported an average SD_W_ of 0.006-0.010 ppm. We observed a within-site SD_w_ range of 0.0009-0.004 ppm and cross-site SD_w_ range of 0.001-0.005 ppm at 7T. Compared to 3T studies, this is a 2.0-5.3 fold decrease in the within-site SD_w_, and a 2.1-4.8 decrease in the cross-site SD_w_., and a 2.1-4.8 decrease in the cross-site SD_w_.

The range of within-site SD_w_ values for R_2_* was averaged 0.0003-0.001 ms^-1^ in our study and the cross-site SD_w_ range was 0.0005-0.001 ms^-1^. The cross-site values are comparable to the *same site* reported at 3T: 0.0005-0.0009 ms^-1^ (Feng et al., 2018), 0.0006-0.002 ms^-1^ (Santin et al., 2017). Compared to the latter, our cross-site results show a 1.1-3.4 improvement over the same brain regions in R2* variability.

The study from Hinoda et al. (2015) measured QSM reproducibility at 1.5T and 3T by scanning subjects twice on each of the scanners. They showed there is a 1.1-2.1 fold decrease in the upper and lower limits in Bland-Altman plots at 3 T compared to 1.5 T, which is in line with the expected signal-to-noise ratio (SNR) increase between these two field strengths (Edelstein et al., 1986; Wardlaw et al., 2012). Compared to 3T reports, there is, on average, an improvement of approximately 3-fold in our QSM and R_2_* 7T measurements of reproducibility. This is in line with the expected SNR increase in brain imaging from 3T to 7T (Pohmann et al., 2015).

The higher values of cross-site SD_w_ compared to the within-site values in our study may be attributed to the different gradient systems and automatic distortion corrections used in the different scanner platforms and to the different approaches to shimming, which lead to different geometrical distortions and dropout regions (Figure 3 and 4, Supplementary Material 2) (Yang et al., 2010). In our study we verified that not only regions in the cortex close to air-tissue interfaces show differences in B_0_ across scanners, but also large subcortical regions such as the CN, the Pu and the GP ROIs. We also showed that the use of a non-linear registration method (here, “SyN” in ANTs) significantly reduced the inter-scanner variability of cortical QSM compared to rigid-body registration, indicating that differences in geometric distortion across scanners were present. The R_2_* results for both cortical and subcortical structures also show significantly lower inter-scanner variability when a non-linear registration was used. For QSM, higher cross-site variability may also be attributed to the head orientation with respect to B_0_ (Lancione et al., 2017; Li et al., 2017). Our results indicate head orientation varied somewhat between scans and there was greater variation between sites than intra-site; we also observed a consistent negative correlation between χ and head orientation (*θ*). Using a linear model to attempt to regress-out the effects of head rotation improved the reproducibility of both within-site and cross-site data. It also reduced the penalty for multi-site scanning vs single-site scanning, but not completely.

In this study, the reproducibility of QSM using single-echo, high-resolution (0.7 mm isotropic resolution; TE=20ms) and multi-echo standard-resolution (1.4 mm isotropic resolution; TE=4, 9, 14, 19, 24, 29, 34 and 39 ms) protocols were compared, and the results show that the multi-echo QSM data has a significantly higher variability than single-echo QSM. Although multi-echo phase data has been combined with a magnitude-weighted least squares regression of phase to echo time, it may carry inconsistent phase accumulation across echoes that were inconsistently unwrapped. This is also particularly relevant for regions of large field inhomogeneities, where phase accumulation in late echoes could exceed ±π between neighbouring voxels, resulting in multiple phase wraps, which the unwrapping algorithm maybe unable to correct (Cronin et al., 2017). This has also been verified on the analysis of QSM data from the cortical ROIs reconstructed with different numbers of echoes: long echo times increase significantly the test-retest variability. Alternative phase unwrapping methods exist such as to perform temporal phase unwrapping across all echo times on the multi-echo data (Liu et al., 2013; Schweser et al., 2013).

It has been shown that resolution influences QSM estimation. Haacke et al. (2015) showed on phantom data that by decreasing slice thickness from 3 mm to 0.5 mm partial volume effects are reduced, absolute susceptibility values decrease, and accuracy improves up to 25%. Similar findings on in vivo brain data are reported in Sun et al. (2017) (single-echo data) and Karsa et al. (2018) (multi-echo data). Our results support the suggestion that a reduction of partial volume effects at higher-resolution might play a role in decreasing both test-retest and cross-site variability on the single-echo high-resolution data compared to the multi-echo low-resolution data.

R_2_* values show significantly lower variability, reflected in the higher ICC within and across-sites compared to corresponding values for χ in subcortical areas. This may be because the χ estimation is globally more sensitive to background field inhomogeneity compared to magnitude data. However, in orbitofrontal and lower temporal regions large through-plane field variations from tissue-air interfaces dominate the field changes and produce dropouts in the signal magnitude and increase the background phase, affecting both QSM and R_2_* maps by increasing variability and decreasing ICC across sites. In addition, because of large field variations, the estimated cortical R_2_* increases significantly when late echo times are used for the fitting, but this effect is not seen in subcortical areas.

QSM can only determine relative susceptibility differences (Cheng et al., 2009) and most approaches to calculation of susceptibility from measured phase yield maps in which the average value of susceptibility is zero over the masked imaging volume. Issues related to referencing of QSM data have been investigated (Feng et al., 2018; Straub et al., 2017), with aim of finding a reference region or tissue to which all susceptibility values are referred that produces well-defined and reproducible values of susceptibility. Here we investigated how the choice of reference affects the within-site and cross-site variability of measured susceptibility at ultra-high-field. We tested three accepted reference regions: total whole brain signal, “wb”, whole brain CSF eroded in order to exclude any pial or skull surfaces, “csf”, and a manually selected cylindrical ROI in the right ventricle, “cyl”. We found that the “cyl” referencing generally increased the variability of the cross-site and within-site susceptibility measurements in cortical and subcortical ROIs compared to “wb” referencing. In the case of the multi-echo acquisition the “csf” referencing also increased the variability relative to “wb” data. This may be because of imprecision in systematically obtaining average QSM signal from CSF regions. Referencing using a small ROI in the ventricles might be prone to subjectivity given the natural variation in ventricle size in healthy subjects and in disease. Furthermore, the ventricles do not contain pure CSF: they are traversed by blood vessels with a different χ (Sullivan et al., 2002). This makes whole-brain referencing attractive in many situations. Yet, in patient cohorts where there is substantial iron load in subcortical structures (Snyder and Connor, 2009), whole brain referencing might not be an appropriate approach. In this case, the more appropriate approach will be to choose a small reference region which shows no changes in the particular disease to be “zero” susceptibility at a cost of a slight increase in SD.

To eliminate operator-dependent bias in segmentation when determining brain structures, we have analysed data using both manual and atlas-based segmentation. From our results, manual ROIs showed significantly lower variability compared to atlas-based methods. This happens because of imprecision in registration between MNI and subject space as well as the empirical thresholding that was chosen to obtain the subcortical ROIs. This resulted in larger ROIs being derived from the atlas-based method compared to the manual method (Wilcoxon test, CN: p=0.014; Pu: p=0.00018; GP: p=0.0010). Overestimation of the region (Figure 5, Supplementary Material 2) meant including boundary voxels that, generally, have lower susceptibility (white-matter, for example), lowering the average χ and R_2_*. However, traditional manual drawing of ROIs for cohort studies is difficult, time consuming and potentially unsuitable as it biases results towards particular cohorts (Collins et al., 2003) so it may not always be the most appropriate approach.

In this study, harmonized protocols were produced for all five scanners without any significant sequence alterations, as a product 3D gradient echo (GE) sequence was readily available on all systems (the product ‘gre’ sequence from Siemens and the product ‘ffe’ from Philips). The protocols and an example dataset are provided in (Clarke, 2018). Generally, we also relied on the vendors’ reconstruction. However, at the end of the reconstruction pipeline of the Siemens systems we adopted a different coil combination approach based on Roemer et al. (1990) and Walsh et al. (2000), to match the SENSE approach implemented on Philips scanners (Pruessmann et al., 1999; Robinson et al., 2017). This was required due to artifacts appearing on phase images in Siemens data reconstructed with the vendor’s pipeline, such as open-ended fringe lines or singularities (Chavez et al., 2002) (Figure 2, Supplementary Material 2). These reduce the consistency of the QSM results (Santin et al., 2017). However, other coil combination methods such as a selective channel combination approach (Vegh et al., 2016) or the COMPOSER (COMbining Phase data using a Short Echo-time Reference scan) method (Bollmann et al., 2018) have also been shown to reduce open-ended fringe lines and noise in the signal phase. For future investigations, the raw k-space data collected from all sites in this study has been stored and is available from the authors upon request.

On the QSM reconstruction, an imperfect background field filtering can influence the reproducibility of QSM data. For this reason, we performed background removal in two steps as implemented in QSMbox v2.0 and as described in (Acosta-Cabronero et al., 2018): first with the LBV approach and then followed by the vSMV method. Regularized field-to-susceptibility inversion strategies have been proposed to overcome the ill-posed problem in QSM with data acquired at a single head orientation (de Rochefort et al., 2010). We opted to use the MSDI implementation in QSMbox v2.0 (Acosta-Cabronero et al., 2018), as it ranked top-10 in all metrics of the 2016 QSM Reconstruction Challenge (Langkammer et al., 2018), and also now includes a new self-optimized local scale, which results in a better preservation of phase noise texture and low susceptibility contrast features. On the second step, the regularization factor, λ, used for this study was set to 10^2.7^, as recommended by Acosta-Cabronero et al. (2018) based on an L-curve analysis (Hansen et al., 1993) with high-resolution 7T data.

The standard multi-echo GE protocol in this study was produced as a harmonised sequence that could be performed at all sites, with a relatively short acquisition time (approximately 5 minutes), which is acceptable for patient studies. Mid-brain structures such as the basal ganglia are identifiable, yet small subcortical structures will suffer from partial-volume effects, which could be a limitation of this harmonized protocol for future ultra-high field multi-site studies.

At ultra-high field there can be variations in SNR in magnitude data caused by the variable B_1_^+^ across the brain (Abduljalil et al., 2003). As R_2_* is estimated voxel-wise, and as there is always a reasonable SNR on the magnitude data, the coefficient in the exponential fit that estimates R_2_* will not be strongly affected by variations in B_1_^+^. QSM maps are estimated from filtered phase data which is not strongly affected by transmit B_1_ variations. On our data, no correlations were found between QSM or R_2_* maps and B_1_ flip-angle maps collected in the same session (Figure 6, Supplementary Material 2).

To minimise confounding effects of age or pathology, we assessed test-retest reliability and cross-site variability with ten healthy young subjects. The cross-site, between-subject standard-deviation, SD_b_, measured in this study was evaluated together with healthy and Parkinson’s disease data from (Langkammer et al., 2016). A power analysis revealed a sample size that would have been required for a multi-site clinical study in each ROI as shown in Figure 8. For all the significant ROIs the number of subjects that would have been required per group was less or equal to 44. Since this is lower than the sample size we have used in this study (90 healthy volunteer scans) and the numbers in the Langkammer study (66 patients and 58 control subjects), it gives strong confidence of feasibility for future 7T QSM clinical studies.

**Figure 8.**
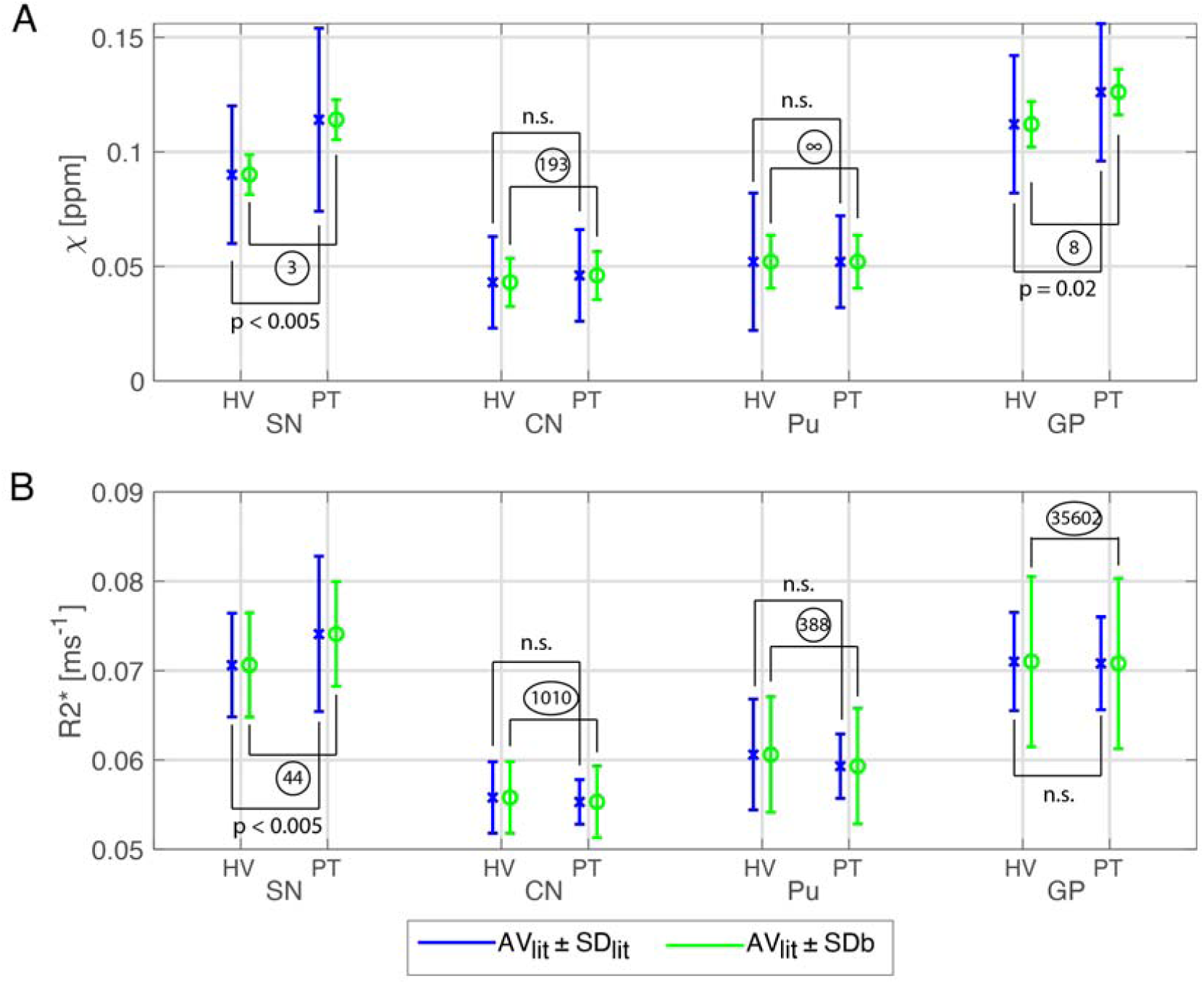
Illustration of the feasibility of a 7T QSM clinical study. χ (A) and R_2_* (B) for four ROIs (Substantia Nigra, SN; Caudate Nucleus, CN; Putamen, Pu; Globus Pallidus, GP) from healthy volunteer (HV) and synthetic “patient” (PT) data for which AV_lit_ and SD_lit_ were obtained from Langkammer et al. (2016) and SD_b_ were calculated from data of the current study. AV_lit_ values for R_2_* were linearly scaled to 7T according to Yao et al. (2007). Blue bars show the AV_lit_ ± SD_lit_ and green bars the AV_lit_ ± SD_b_. Statistical differences between HV and PT obtained from Langkammer et al. (2016) are also shown. For each ROI, the sample size that would have been needed to give a significant effect was calculated from the group means, AV_lit_, and the SD_b_ per ROI and is shown in circles. Multi-echo χ-maps were calculated with data from all eight echoes.

## 5. Conclusion

We investigated test-retest reliability and reproducibility of T_2_*-weighted imaging protocols at ultra-high field MRI. Considering the increase in susceptibility effects at 7T, we found that variability of measurements of QSM χ and R_2_* in the basal ganglia are reduced compared to reports from lower field strengths, 1.5T and 3T. Scanner hardware differences give more modest improvements for cortical measurements of QSM χ and R_2_*. Multi-echo protocols do not benefit from long echo times as these increase the imprecision in the estimation of QSM. We suggest that 7T MRI is suitable for multicentre quantitative analyses of brain iron, in health and disease.

## 6. Acknowledgements

The UK7T Network and this work was funded by the UK’s Medical Research Council (MRC) [MR/N008537/1]. We thank Dr. Julio Acosta-Cabronero for making the QSMbox publically available and Prof. David Porter for the support in the UK7T network.

## 7. Centre funding

The Wellcome Centre for Integrative Neuroimaging is supported by core funding from the Wellcome Trust (203139/Z/16/Z).

Cardiff University Brain Research Imaging Centre is supported by the UK Medical Research Council (MR/M008932/1) and the Wellcome Trust (WT104943).

This research was co-funded by the NIHR Cambridge Biomedical Research Centre. The views expressed are those of the author(s) and not necessarily those of the NHS, the NIHR or the Department of Health and Social Care. The Cambridge 7T MRI facility is co-funded by the University of Cambridge and the Medical Research Council (MR/M008983/1).

## 8. Individual funding

CTR is funded by a Sir Henry Dale Fellowship from the Wellcome Trust and the Royal Society [098436/Z/12/B]. JBR is supported by the Wellcome Trust (WT103838).

## References

Abdul-Rahman, H., Gdeisat, M., Burton, D., Michael, L., 2005. Fast three-dimensional phase-unwrapping algorithm based on sorting by reliability following a non-continuous path. App. Optic. 46, 6623–6635. doi: 10.1117/12.611415

Abduljalil, A.M., Schmalbrock, P., Novak, V., Chakeres, D.W. 2003. Enhanced gray and white matter contrast of phase susceptibility-weighted images in ultra-high-field magnetic resonance imaging. Journal of Magnetic Resonance Imaging: An Official Journal of the International Society for Magnetic Resonance in Medicine. 18(3), 284–90. doi: 10.1002/jmri.10362

Acosta-Cabronero, J., Milovic, C., Mattern, H., Tejos, C., Speck, O., Callaghan, M.F., 2018. A robust multi-scale approach to quantitative susceptibility mapping. NeuroImage 183, 7–24. doi: 10.1016/j.neuroimage.2018.07.065

Acosta-Cabronero, J., Betts, M.J., Cardenas-Blanco, A., Yang, S., Nestor, P.J., 2016. In Vivo MRI mapping of brain iron deposition across the adult lifespan. J. Neurosci. 36, 364–374. doi: 10.1523/JNEUROSCI.1907-15.2016

Acosta-Cabronero, J., Williams, G.B., Cardenas-Blanco, A., Arnold, R.J., Lupson, V., Nestor, P.J., 2013. In vivo quantitative susceptibility mapping (QSM) in Alzheimer’s disease. PLoS One 8, e81093. doi: 10.1371/journal.pone.0081093

Barbosa, J.H.O., Santos, A.C., Salmon, C.E.G., 2015. Susceptibility weighted imaging: differentiating between calcification and hemosiderin. Radiologia brasileira 48(2), 93–100. doi: 10.1590/0100-3984.2014.0010

Betts, M.J., Acosta-Cabronero, J., Cardenas-Blanco, A., Nestor, P.J., Düzel, E., 2016. High-resolution characterisation of the aging brain using simultaneous quantitative susceptibility mapping (QSM) and R2* measurements at 7 T. Neuroimage 138, pp.43–63. doi: 10.1016/j.neuroimage.2016.05.024

Bilgic, B., Pfefferbaum, A., Rohlfing, T., Sullivan, E.V., Adalsteinsson, E., 2012. MRI estimates of brain iron concentration in normal aging using quantitative susceptibility mapping. Neuroimage 59(3), 2625–2635. doi: 10.1016/j.neuroimage.2011.08.077

Blazejewska, A.I., Al-Radaideh, A.M., Wharton, S., Lim, S.Y., Bowtell, R.W., Constantinescu, C.S., Gowland, P.A., 2015. Increase in the iron content of the substantia nigra and red nucleus in multiple sclerosis and clinically isolated syndrome: a 7 Tesla MRI study. J. Magn. Reson. Imaging 41, 1065–1070. doi: 10.1002/jmri.24644

Bollmann, S., Robinson, S.D., O’Brien, K., Vegh, V., Janke, A., Marstaller, L., Reutens, D., Barth, M., 2018. The challenge of bias-free coil combination for quantitative susceptibility mapping at ultra-high field. Magn. Reson. Med. 79(1), 97–107. doi: 10.1002/mrm.26644

Chavez, S., Xiang, Q., Li, A., 2002. Understanding phase maps in MRI: a new cutline phase unwrapping method. IEEE Trans Med Imaging 21, 966–977. doi: 10.1109/TMI.2002.803106

Cheng, Y.-C.N., Neelavalli, J., Haacke, E.M., 2009. Limitations of calculating field distributions and magnetic susceptibilities in MRI using a Fourier based method. Phys. Med. Biol. 54(5), 1169–1189. doi: 10.1088/0031-9155/54/5/005

Choi, S., Li, X., Harrison, D.M., 2019. The impact of coregitration of gradient recalled echo images on quantitative susceptibility and R2* mapping at 7T. bioRxiv. doi: 10.1101/529891

Clarke, W.T., 2018. UK7T Network harmonized neuroimaging protocols. https://ora.ox.ac.uk/objects/uuid:55ca977f-62df-4cbf-b300-2dc893e36647.

Clarke, W.T., Mougin, O., Driver, I.D., Rua, C., Morgan, A., Asghar, M., Clare, S., Francis, S., Wise, R., Rodgers, C.T., Carpenter, T.A., Muir, K., Bowtell, R., 2019. Multi-site harmonization of 7 Tesla MRI neuroimaging protocols. NeuroImage 206, 116335. doi: 10.1016/j.neuroimage.2019.116335

Cobzas, D., Sun, H.F., Walsh, A.J., Lebel, R.M., Blevins, G., Wilman, A.H., 2015. Subcortical gray matter segmentation and voxel-based analysis using transverse relaxation and quantitative susceptibility mapping with application to multiple sclerosis. J. Magn. Reson. Imaging 42(6), 1601–1610. doi: 10.1002/jmri.24951

Collins, D.L., Zijdenbos, A.P., Paus, T., Evans, A.C., 2003. Use of registration for cohort studies. Medical image registration.

de Rochefort, L., Liu, T., Kressler, B., Liu, J., Spincemaille, P., Lebon, V., Wu, J., Wang, Y., 2010. Quantitative susceptibility map reconstruction from MR phase data using Bayesian regularization: validation and application to brain imaging. Magn. Reson. Med. 63, 194–206. doi: 10.1002/mrm.22187

Deh, K., Nguyen, T.D., Eskreis-Winkler, S., Prince, M.R., Spincemaille, P., Gauthier, S., Kovanlikaya, I., Zhang, Y., Wang, Y., 2015. Reproducibility of quantitative susceptibility mapping in the brain at two field strengths from two vendors. J. Magn. Reson. Imaging 42, 1592–1600. doi: 10.1002/jmri.24943

Deistung, A., Schäfer, A., Schweser, F., Biedermann, U., Güllmar, D., Trampel, R., Turner, R., Reichenbach, J.R., 2013. High-resolution MR imaging of the human brainstem in vivo at 7 Tesla. Frontiers in human neuroscience 7, 710. doi: 10.3389/fnhum.2013.00710

Düzel, E., Acosta-Cabronero, J., Berron, D., Biessels, G.J., Björkman-Burtscher, I., Bottlaender, M., Bowtell, R., Buchem, M.V., Cardenas-Blanco, A., Boumezbeur, F., Chan, D., 2019. European Ultrahigh-Field Imaging Network for Neurodegenerative Diseases (EUFIND). Alzheimer’s & Dementia: Diagnosis, Assessment & Disease Monitoring, 11(1), 538–549. doi: 10.1016/j.dadm.2019.04.010

Duyn, J.H., van Gelderen, P., Li, T.Q., de Zwart, J.A., Koretsky, A.P., Fukunaga, M., 2007. High-field MRI of brain cortical substructure based on signal phase. Proceedings of the National Academy of Sciences, 104(28), 11796–11801. doi: 10.1073/pnas.0610821104

Edelstein, W.A., Glover, G.H., Hardy, C.J., Redington, R.W., 1986. The intrinsic signal-to-noise ratio in NMR imaging. Magnetic resonance in medicine, 3(4), 604–618. doi: 10.1002/mrm.1910030413

Ehses, P., Brenner, D., Stirnberg, R., Pracht, E.D., Stöcker, T., 2019. Whole-brain B1-mapping using three-dimensional DREAM. Magn. Reson. Med. 82(3), 924–934. doi: 10.1002/mrm.27773

Eskreis-Winkler, S., Zhang, Y., Zhang, J., Liu, Z., Dimov, A., Gupta, A., Wang, Y., 2017. The clinical utility of QSM: disease diagnosis, medical management, and surgical planning. NMR in Biomedicine 30(4), p.e3668. doi: 10.1002/nbm.3668

Feng, X., Deistung, A., Reichenbach, J.R., 2018. Quantitative susceptibility mapping (QSM) and R2* in the human brain at 3 T: Evaluation of intra-scanner repeatability. Z. Med. Phys. 28, 36–48. doi: 10.1016/j.zemedi.2017.05.003

Gelman, N., Gorell, J.M., Barker, P.B., Savage, R.M., Spickler, E.M., Windham, J.P., Knight, R.A., 1999. MR imaging of human brain at 3.0 T: preliminary report on transverse re-laxation rates and relation to estimated iron content. Radiology 210, 759–767. doi: 10.1148/radiology.210.3.r99fe41759

Haacke, E.M., Cheng, N., House, M.J., Liu, Q., Neelavalli, J., Ogg, R.J., Khan, A., Ayaz, M., Kirsch, W., Obenaus, A., 2005. Imaging iron stores in the brain using magnetic resonance imaging. Magn. Reson. Imag. 23, 1–25. doi: 10.1016/j.mri.2004.10.001

Haacke, E.M., Liu, S., Buch, S., Zheng, W., Wu, D., Ye, Y., 2015. Quantitative susceptibility mapping: current status and future directions. Magnetic resonance imaging, 33(1), 1–25. doi: 10.1016/j.mri.2014.09.004

Hansen, P.C., O’Leary, D.P., 1993. The use of the l-curve in the regularization of discrete ill-posed problems. SIAM J Sci Comput 14(6), 1487–1503. doi: 10.1137/0914086

He, X., Yablonskiy, D.A., 2009. Biophysical mechanisms of phase contrast in gradient echo MRI. Proc. Natl. Acad. Sci. U.S.A. 106, 13558–13563. https://doi.org/10.1073/pnas.0904899106

Hinoda, T., Fushimi, Y., Okada, T., Fujimoto, K., Liu, C., Yamamoto, A., Okada, T., Kido, A., Togashi, K., 2015. Quantitative susceptibility mapping at 3 T and 1.5 T: evaluation of consistency and reproducibility. Invest. Radiol. 50, 522–530. doi: 10.1097/RLI.0000000000000159

House, M.J., Pierre, T.G.S., Kowdley, K.V., Montine, T., Connor, J., Beard, J., Berger, J., Siddaiah, N., Shankland, E., Jin, L.W., 2007. Correlation of proton transverse relaxation rates (R2) with iron concentrations in postmortem brain tissue from Alzheimer’s disease patients. Magn. Reson. Med. 57, 172–180. doi: 10.1002/mrm.21118

Karsa, A., Punwani, S., Shmueli, K., 2019. The effect of low resolution and coverage on the accuracy of susceptibility mapping. Magnetic resonance in medicine, 81(3), 1833–1848. doi: 10.1002/mrm.27542

Keuken, M.C., Bazin, P.L., Backhouse, K., Beekhuizen, S., Himmer, L., Kandola, A., Lafeber, J.J., Prochazkova, L., Trutti, A., Schäfer, A., Turner, R., 2017. Effects of aging on T1, T2*, and QSM MRI values in the subcortex. Brain Structure and Function 222(6), 2487–2505. doi: 10.1007/s00429-016-1352-4

Lancione, M., Tosetti, M., Donatelli, G., Cosottini, M., Costagli, M., 2017. The impact of white matter fiber orientation in single-acquisition quantitative susceptibility mapping. NMR Biomed. 30, e3798. doi: 10.1002/nbm.3798

Langkammer, C., Schweser, F., Shmueli, K., Kames, C., Li, X., Guo, L., Milovic, C., Kim, J., Wei, H., Bredies, K., Buch, S., 2018. Quantitative susceptibility mapping: report from the 2016 reconstruction challenge. Magn. Reson. Med. 79(3), 1661–73. doi: 10.1002/mrm.26830

Langkammer, C., Pirpamer, L., Seiler, S., Deistung, A., Schweser, F., Franthal, S., Homayoon, N., Katschnig-Winter, P., Koegl-Wallner, M., Pendl, T., Stoegerer, E.M., 2016. Quantitative susceptibility mapping in Parkinson’s disease. PLoS One, 11(9), e0162460. doi: 10.1371/journal.pone.0162460

Langkammer, C., Schweser, F., Krebs, N., Deistung, A., Goessler, W., Scheurer, E., Sommer, K., Reishofer, G., Yen, K., Fazekas, F., Ropele, S., Reichenbach, J.R., 2012. Quantitative susceptibility mapping (QSM) as a means to measure brain iron? A post mortem validation study. Neuroimage 62(3), 1593–1599. doi: 10.1016/j.neuroimage.2012.05.049

Langkammer, C., Krebs, N., Goessler, W., Scheurer, E., Ebner, F., Yen, K., Fazekas, F., Ropele, S., 2010. Quantitative MR imaging of brain iron: a postmortem validation study. Radiology 257(2), 455–462. doi: 10.1148/radiol.10100495

Lee, J., Shmueli, K., Kang, B.T., Yao, B., Fukunaga, M., van Gelderen, P., Palumbo, S., Bosetti, F., Silva, A.C., Duyn, J.H., 2012. The contribution of myelin to magnetic susceptibility-weighted contrasts in high-field MRI of the brain. Neuroimage 59, 3967–3975. doi: 10.1016/j.neuroimage.2011.10.076

Li, G., Zhai, G., Zhao, X., An, H., Spincemaille, P., Gillen, K.M., Ku, Y., Wang, Y., Huang, D., Li, J., 2019. 3D texture analysis within substantia nigra of Parkinson’s disease patients on quantitative susceptibility maps and R2* maps. NeuroImage 188, 465–472. doi: 10.1016/j.neuroimage.2018.12.041

Li, L., Leigh, J.S., 2004. Quantifying arbitrary magnetic susceptibility distributions with MR. Magn. Reson. Med. 51, 1077–1082. doi: 10.1002/mrm.20054

Li, W., Liu, C., Duong, T.Q., van Zijl, P.C., Li, X., 2017. Susceptibility tensor imaging (STI) of the brain. NMR Biomed. 30(4), p.e3540. doi: 10.1002/nbm.3540

Lin, P.Y., Chao, T.C., Wu, M.L., 2015. Quantitative susceptibility mapping of human brain at 3T: a multisite reproducibility study. AJNR Am J. Neuroradiol. 36, 467–474. doi: 10.3174/ajnr.A4137

Liu, T., Wisnieff, C., Lou, M., Chen, W., Spincemaille, P., Wang, Y., 2013. Nonlinear formulation of the magnetic field to source relationship for robust quantitative susceptibility mapping. Magn Reson Med. 69(2), 467–76. doi: 10.1002/mrm.24272

Lotfipour, A.K., Wharton, S., Schwarz, S.T., Gontu, V., Schaefer, A., Peters, A.M., Bowtell, R.W., Auer, D.P., Gowland, P.A., Bajaj, P.S., 2012. High resolution magnetic susceptibility mapping of the substantia nigra in Parkinson’s disease. J. Magn. Reson. Imag. 35, 48–55. doi: 10.1002/jmri.22752

Makhlouf, S.A., Parker, F.T., Berkowitz, A.E., 1997. Magnetic hysteresis anomalies in ferritin. Physical Review B, 55(22), R14 717–R14 720. doi: 10.1103/PhysRevB.55.R14717

Moeller, H.E., Bossoni, L., Connor, J.R., Crichton, R.R., Does, M.D., Ward, R.J., Zecca, L., Zucca, F.A., Ronen, I., 2019. Iron, myelin, and the brain: Neuroimaging meets neurobiology. Trends in neurosciences, 42(6), 384–401. doi: 10.1016/j.tins.2019.03.009

Mougin, O., Clarke, W., Driver, I., Rua, C., Morgan, A.T., Francis, S., Muir, K., Carpenter, A., Rodgers, C., Wise, R., Porter, D., Clare, S., Bowtell, R., 2019. Robustness of PSIR segmentation and R1 mapping at 7T: a travelling head study. Proc. Intr. Soc. Mag. Reson. Med. 27, 237.

Nehrke, K., Bornert, P., 2012. DREAM--a novel approach for robust, ultrafast, multislice B(1) mapping. Magn. Reson. Med. 68(5), 1517–1526. doi: 10.1002/mrm.24158

Pei, M., Nguyen, T.D., Thimmappa, N.D., Salustri, C., Dong, F., Cooper, M.A., Li, J., Prince, M.R., Wang, Y., 2015. Algorithm for fast monoexponential fitting based on auto-regression on linear operations (ARLO) of data. Magn. Reson. Med. 73, 843–850. doi: 10.1002/mrm.25137

Pohmann, R., Speck, O., Scheffler, K., 2016. Signal-to-noise ratio and MR tissue parameters in human brain imaging at 3, 7, and 9.4 tesla using current receive coil arrays. Magnetic resonance in medicine, 75(2), 801–809. doi: 10.1002/mrm.25677

Pruessmann, K.P., Weiger, M., Scheidegger, M.B., Boesiger, P., 1999. SENSE: sensitivity encoding for fast MRI. Magn. Reson. Med. 42(5), 952–962. doi: 10.1002/(SICI)1522-2594(199911)42:5<952::AID-MRM16>3.0.CO;2-S

R Core team, 2013. R: A language and environment for statistical computing. R Foundation for Statistical Computing, Vienna, Austria. http://www.R-project.org/.

Reichenbach, J.R., 2012. The future of susceptibility contrast for assessment of anatomy and function. Neuroimage 62, 1311–1315. doi: 10.1016/j.neuroimage.2012.01.004

Reichenbach, J.R., Jonetz-Mentzel, L., Fitzek, C., Haacke, E.M., Kido, D.K., Lee, B.C., Kaiser, W.A., 2001. High-resolution blood oxygen-level dependent MR venography (HRBV): a new technique. Neuroradiology 43, 364–369. doi: 10.1007/s002340000503

Robinson, S.D., Bredies, K., Khabiova, D., Dymerska, B., Marques, J.P., Schweser, F., 2017. An illustrated comparison of processing methods for MR phase imaging and QSM: combining array coil signals and phase unwrapping. NMR Biomed. 30, e3601. doi: 10.1002/nbm.3601

Roemer, P.B., Edelstein, W.A., Hayes, C.E., Souza, S.P., Mueller, O.M., 1990. The NMR phased array. Magn. Reson. Med. 16(2), 192–225. doi: 10.1002/mrm.1910160203

Santin, M.D., Didier, M., Valabregue, R., Yahia Cherif, L., García-Lorenzo, D., Loureiro de Sousa, P., Bardinet, E., Lehéricy, S., 2017. Reproducibility of R2* and quantitative susceptibility mapping (QSM) reconstruction methods in the basal ganglia of healthy subjects. NMR Biomed. 30(4), e3491. doi: 10.1002/nbm.3491

Schweser, F., Deistung, A., Lehr, B.W., Reichenbach, J.R., 2011. Quantitative imaging of intrinsic magnetic tissue properties using MRI signal phase: an approach to in vivo brain iron metabolism?. Neuroimage 54(4), 2789–2807. doi: 10.1016/j.neuroimage.2010.10.070

Schweser. F., Deistung, A., Sommer, K., Reichenbach, J.R., 2013. Toward online reconstruction of quantitative susceptibility maps: superfast dipole inversion. Magn Reson Med. 69(6), 1582–94. doi: 10.1002/mrm.24405

Smith, S.M., 2002. Fast robust automated brain extraction. Human Brain Mapping 17(3), 143–155. doi: 10.1002/hbm.10062

Snyder A.M., Connor J.R., 2009. Iron, the substantia nigra and related neurological disorders. Biochimica et Biophysics Acta 1790, 606–614. doi: 10.1016/j.bbagen.2008.08.005

Spincemaille, P., Liu, Z., Zhang, S., Kovanlikaya, I., Ippoliti, M., Makowski, M., Watts, R., de Rochefort, L., Venkatraman, V., Desmond, P., Santin, M.D., 2019. Clinical integration of automated processing for brain quantitative susceptibility mapping: multi-site reproducibility and single-site robustness. Journal of Neuroimaging 29(6), 689–698. doi: 10.1111/jon.12658

Straub, S., Schneider, T.M., Emmerich, J., Freitag, M.T., Ziener, C.H., Schlemmer, H.P., Ladd, M.E., Laun, F.B., 2017. Suitable reference tissues for quantitative susceptibility mapping of the brain. Magn. Reson. Med. 78, 204–214. doi: 10.1002/mrm.26369

Sullivan, E.V., Pfefferbaum, A., Adalsteinsson, E., Swan, G.E., Carmelli, D., 2002. Differential rates of regional brain change in callosal and ventricular size: a 4-year longitudinal MRI study of elderly men. Cereb. Cortex 12(4), 438–45. doi: 10.1093/cercor/12.4.438

Sun, H., Seres, P., Wilman, A.H., 2017. Structural and functional quantitative susceptibility mapping from standard fMRI studies. NMR in Biomedicine, 30(4), e3619. doi: 10.1002/nbm.3619

Tie-Qiang, T., Gelderen, P., Merkle, H., Talagala, L., Koretsky, A.P., Duyn, J., 2006. Extensive heterogeneity in white matter intensity in high-resolution T2*-weighted MRI of the human brain at 7.0 T. NeuroImage 32, 1032–1040. doi: 10.1016/j.neuroimage.2006.05.053

Vegh, V., O’Brien, K., Barth, M., Reutens, D.C., 2016. Selective channel combination of MRI signal phase. Magn. Reson. Med. 76(5), 1469–1477. doi: 10.1002/mrm.26057

Walsh, D.O., Gmitro, A.F., Marcellin, M.W., 2000. Adaptive reconstruction of phased array MR imagery. Magn. Reson. Med. 43(5), 682–690. doi: 10.1002/(SICI)1522-2594(200005)43:5<682::AID-MRM10>3.0.CO;2-G

Wang, Y., Liu, T., 2015. Quantitative susceptibility mapping (QSM): decoding MRI data for a tissue magnetic biomarker. Magn. Reson. Med. 7, 82–101. doi: 10.1002/mrm.25358

Wardlaw, J.M., Brindle, W., Casado, A.M., Shuler, K., Henderson, M., Thomas, B., Macfarlane, J., Maniega, S.M., Lymer, K., Morris, Z., Pernet, C., 2012. A systematic review of the utility of 1.5 versus 3 Tesla magnetic resonance brain imaging in clinical practice and research. European radiology, 22(11), 2295–2303. doi: 10.1007/s00330-012-2500-8

Weir, J. P., 2005. Quantifying test-retest reliability using the intraclass correlation coefficient and the SEM. J. Strength Cond. Res. 19(1), 231–240.

Wharton, S., Bowtell, R., 2010. Whole-brain susceptibility mapping at high field: a comparison of multiple-and single-orientation methods. Neuroimage 53(2), 515–525. doi: 10.1016/j.neuroimage.2010.06.070

Yacoub, E., Shmuel, A., Pfeuffer, J., Van de Moortele, P.F., Adriany, G., Andersen, P., Vaughan, J.T., Merkle, H., Ugurbil, K., Hu, X., 2001. Imaging brain function in humans at 7 Tesla. Magn. Reson. Med. 45, 588–594. doi: 10.1002/mrm.1080

Yang, X., Sammet, S., Schmalbrock, P., Knopp, M. V., 2010. Postprocessing correction for distortions in T2* decay caused by quadratic cross-slice b0 inhomogeneity. Magn. Reson. Med. 63(1): 1258–1268. doi: 10.1002/mrm.22316

Yao, B., Li, T., van Gelderen, P., Shmueli, K., De Zwart, J.A., Duyn, J.H., 2009. Neuro image susceptibility contrast in high field MRI of human brain as a function of tissue iron content. Neuroimage 44(4), 1259–66. doi: 10.1016/j.neuroimage.2008.10.029

Yao, B., van Gelderen, P., de Zwart, J.A., Duyn, J.H., 2007. Brain iron in MR imaging: R2* and phase shift at different field strengths. Proc. Intr. Soc. Mag. Reson. Med. 15, 2165.

Zhou, D., Liu, T., Spincemaille, P., Wang, Y., 2014. Background field removal by solving the Laplacian boundary value problem. NMR Biomed. 27(3), 312–319. doi: 10.1002/nbm.3064

