## Supplementary Material 2 for "Multi-centre, multi-vendor reproducibility of 7T QSM and R_2_* in the human brain: results from the UK7T study"

Rua et al.

SUPPLEMENTARY MATERIAL 2

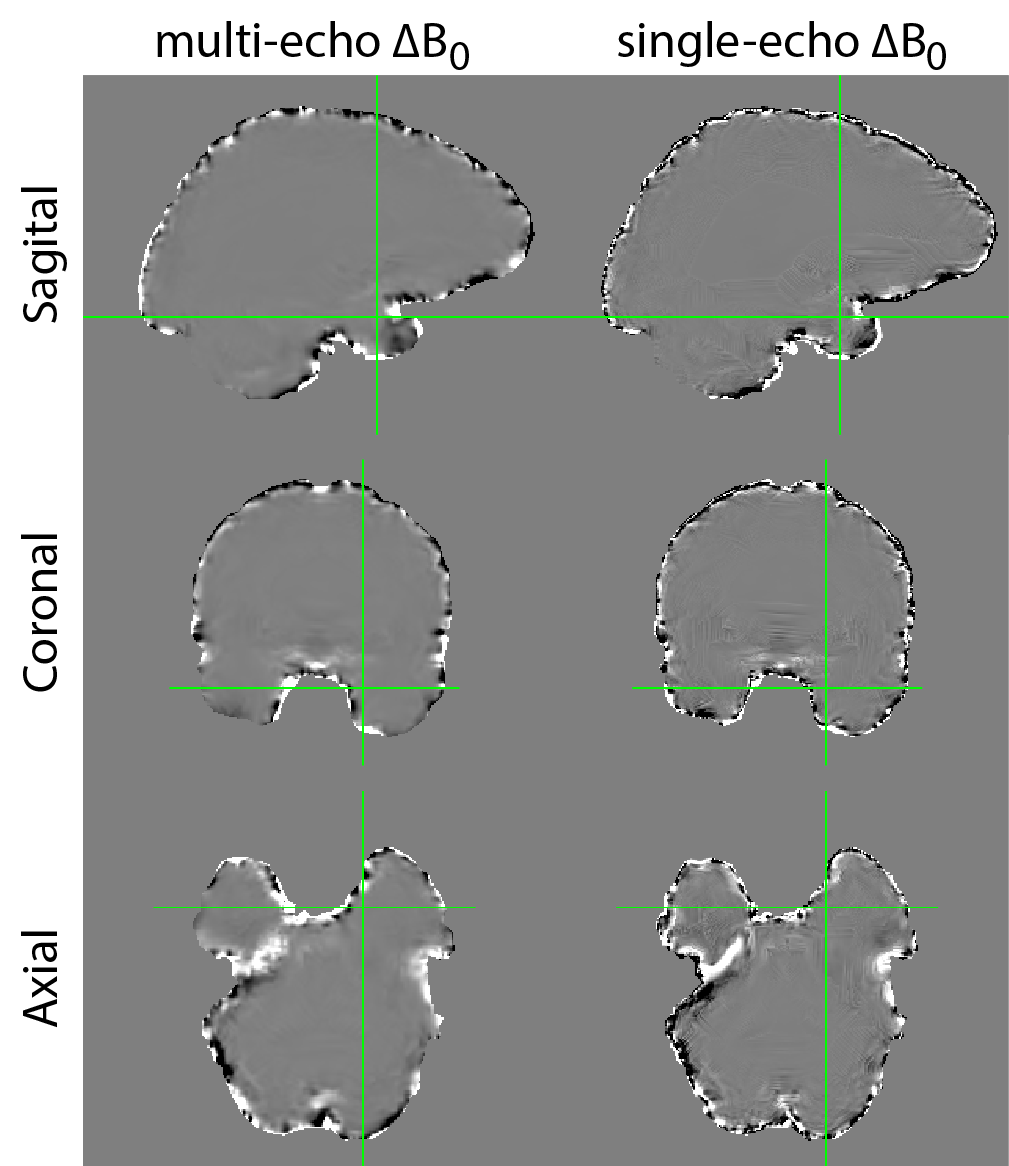

**Figure 1, Supplementary Material 2:** Differences between $\Delta B_{0}$ derived from the multi-echo QSM pipeline (left column) and single-echo QSM pipeline (right column) from an example subject.

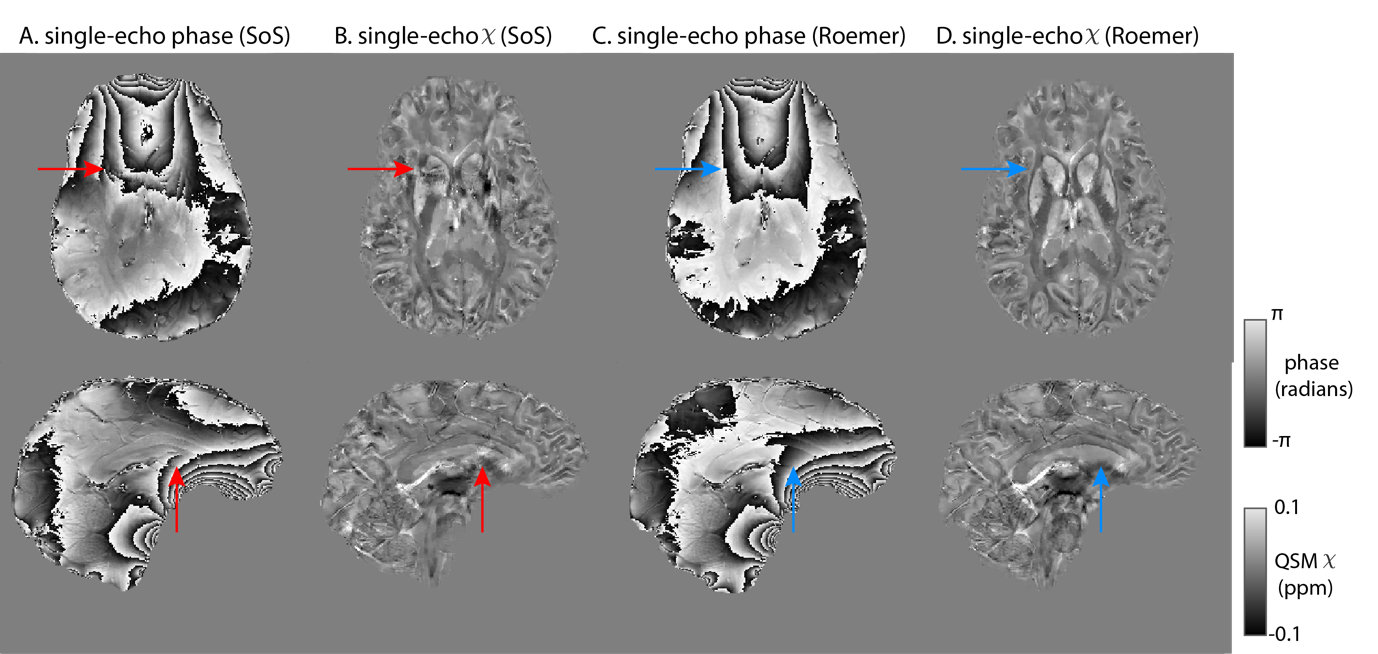
**Figure 2, Supplementary Material 2:** Representative axial and sagittal slices from one example subject. Single-echo T_2_*-w signal phase (TE = 20 ms) using sum-of-squares coil combination in (A) showing singularities in phase data (red arrows) and resulting single-echo χ map in (B). Single-echo T_2_*-w signal phase (TE = 20 ms) using Roemer coil combination in (C) free of open-fringe lines (blue arrows) and corresponding single-echo χ maps (D).

| **Covariate** | **Measure** | **Manual ROIs** | | |
| --- | --- | --- | --- | --- |
|  |  | **Single-echo** $\boldsymbol{\chi}$ | **Multi-echo** $\boldsymbol{\chi}$ | **R_2_*** |
| **Type**  within-site vs cross-site | AV_w_ | 0.053 | 0.65 | **<0.0001** |
|  | SD_w_ | **<0.0001** | **0.033** | **<0.0001** |
|  | ICC | **<0.0001** | **0.017** | **<0.0001** |
| **Registration**  Rigid vs SyN | AV_w_ | **0.0020** | 0.25 | 0.088 |
|  | SD_w_ | 0.72 | 0.51 | **<0.0001** |
|  | ICC | 0.71 | 0.31 | **0.013** |
| **QSM referencing** |  |  |  |  |
| wb vs csf reference | AV_w_ | **<0.0001** | **<0.0001** | - |
|  | SD_w_ | 0.93 | **0.00096** | - |
|  | ICC | 0.75 | 0.71 | - |
| wb vs cyl reference | AV_w_ | **<0.0001** | **<0.0001** | - |
|  | SD_w_ | **<0.0001** | **0.00064** | - |
|  | ICC | 0.099 | 0.46 | - |
| **Multi-Echo T_2_*** |  |  |  |  |
| 1 echo vs 2 echoes | AV_w_ | - | **<0.0001** | - |
|  | SD_w_ | - | **0.029** | - |
|  | ICC | - | 0.27 | - |
| 1 echo vs 5 echoes | AV_w_ | - | **<0.0001** | - |
|  | SD_w_ | - | **0.032** | - |
|  | ICC | - | 0.71 | - |
| 1 echo vs 8 echoes | AV_w_ | - | **<0.0001** | - |
|  | SD_w_ | - | **<0.0001** | - |
|  | ICC | - | **0.009** | - |
| 2 echoes vs 5 echoes | AV_w_ | - | 0.36 | - |
|  | SD_w_ | - | 1.00 | - |
|  | ICC | - | 0.18 | - |
| 2 echoes vs 8 echoes | AV_w_ | - | 0.98 | - |
|  | SD_w_ | - | 0.14 | - |
|  | ICC | - | **0.0011** | - |
| 3 echoes vs 4 echoes | AV_w_ | - | 0.42 | 0.71 |
|  | SD_w_ | - | 0.98 | 0.35 |
|  | ICC | - | 0.86 | 0.65 |
| 3 echoes vs 6 echoes | AV_w_ | - | 0.61 | 0.51 |
|  | SD_w_ | - | 0.11 | 0.060 |
|  | ICC | - | 0.14 | 0.43 |
| 3 echoes vs 8 echoes | AV_w_ | - | 0.75 | 0.58 |
|  | SD_w_ | - | **0.014** | **0.020** |
|  | ICC | - | **0.0017** | 0.49 |
| 4 echoes vs 6 echoes | AV_w_ | - | 0.70 | 0.78 |
|  | SD_w_ | - | 0.13 | 0.34 |
|  | ICC | - | 0.069 | 0.73 |
| 5 echoes vs 7 echoes | AV_w_ | - | 0.54 | 0.99 |
|  | SD_w_ | - | 0.16 | 0.81 |
|  | ICC | - | 0.066 | 0.99 |
| 6 echoes vs 8 echoes | AV_w_ | - | 0.36 | 0.90 |
|  | SD_w_ | - | 0.64 | 0.667 |
|  | ICC | - | **0.027** | 0.95 |
| 7 echoes vs 8 echoes | AV_w_ | - | 0.65 | 0.93 |
|  | SD_w_ | - | 0.97 | 0.85 |
|  | ICC | - | 0.19 | 0.97 |
| **ROI Selection** manual vs atlas-based subcortical ROIs | AV_w_ | **<0.0001** | **<0.0001** | **<0.0001** |
|  | SD_w_ | **<0.0001** | **<0.0001** | **<0.0001** |
|  | ICC | **0.00021** | **0.0023** | **0.012** |

**Table 1, Supplementary Material 2:** p-values obtained from the general linear model on the AVw, SDw and ICC measurements obtained from single-echo QSM, multi-echo QSM and R2* maps in the manual ROIs.

| **Covariate** | **Measure** | **Atlas-based cortical ROIs** | | |
| --- | --- | --- | --- | --- |
|  |  | **Single-echo** $\boldsymbol{\chi}$ | **Multi-echo** $\boldsymbol{\chi}$ | **R_2_*** |
| **Type**  within-site vs cross-site | AVw | 0.23 | 0.9298 | 0.1042 |
|  | SDw | **<0.0001** | **<0.0001** | **<0.0001** |
|  | ICC | **<0.0001** | **<0.0001** | **<0.0001** |
| **Registration**  Rigid vs SyN | AVw | **0.00022** | 0.33 | 0.13 |
|  | SDw | **<0.0001** | 0.12 | **<0.0001** |
|  | ICC | **<0.0001** | **0.034** | **<0.0001** |
| **QSM referencing** |  |  |  |  |
| wb vs csf reference | AVw | **<0.0001** | **<0.0001** | - |
|  | SDw | **<0.0001** | **<0.0001** | - |
|  | ICC | **<0.0001** | **<0.0001** | - |
| wb vs cyl reference | AVw | **<0.0001** | **<0.0001** | - |
|  | SDw | **<0.0001** | **<0.0001** | - |
|  | ICC | **0.021** | 0.22 | - |
| **Multi-Echo T2*** |  |  |  |  |
| 1 echo vs 2 echoes | AV_w_ | - | **<0.0001** | - |
|  | SD_w_ | - | **0.0012** | - |
|  | ICC | - | 0.96 | - |
| 1 echo vs 5 echoes | AV_w_ | - | **0.00018** | - |
|  | SD_w_ | - | **0.022** | - |
|  | ICC | - | 0.47 | - |
| 1 echo vs 8 echoes | AV_w_ | - | 0.21 | - |
|  | SD_w_ | - | **<0.0001** | - |
|  | ICC | - | **0.00068** | - |
| 2 echoes vs 5 echoes | AV_w_ | - | **0.00057** | - |
|  | SD_w_ | - | 0.43 | - |
|  | ICC | - | 0.33 | - |
| 2 echoes vs 8 echoes | AV_w_ | - | **<0.0001** | - |
|  | SD_w_ | - | **0.00085** | - |
|  | ICC | - | **0.00049** | - |
| 3 echoes vs 4 echoes | AV_w_ | - | 0.18 | **0.00034** |
|  | SD_w_ | - | 0.51 | 0.80 |
|  | ICC | - | 0.93 | 0.35 |
| 3 echoes vs 6 echoes | AV_w_ | - | **<0.0001** | **<0.0001** |
|  | SD_w_ | - | **0.00073** | 0.72 |
|  | ICC | - | 0.065 | **0.041** |
| 3 echoes vs 8 echoes | AV_w_ | - | **<0.0001** | **<0.0001** |
|  | SD_w_ | - | **<0.0001** | 0.92 |
|  | ICC | - | **0.00012** | **0.0086** |
| 4 echoes vs 6 echoes | AV_w_ | - | **0.0010** | **<0.0001** |
|  | SD_w_ | - | **0.0057** | 0.91 |
|  | ICC | - | 0.054 | 0.23 |
| 5 echoes vs 7 echoes | AV_w_ | - | **0.011** | **0.00014** |
|  | SD_w_ | - | **0.0010** | 0.99 |
|  | ICC | - | 0.056 | 0.38 |
| 6 echoes vs 8 echoes | AV_w_ | - | 0.12 | **0.0098** |
|  | SD_w_ | - | **0.023** | 0.82 |
|  | ICC | - | **0.0086** | 0.50 |
| 7 echoes vs 8 echoes | AV_w_ | - | 0.45 | 0.31 |
|  | SD_w_ | - | 0.31 | 0.86 |
|  | ICC | - | 0.12 | 0.76 |
| $\boldsymbol{\Delta}\boldsymbol{B}$ | AVw | **<0.0001** | **0.015** | **<0.0001** |
|  | SDw | 0.75 | **0.0011** | **<0.0001** |
|  | ICC | **<0.0001** | **<0.0001** | **<0.0001** |

**Table 2, Supplementary Material 2:** p-values obtained from the general linear model on the AVw, SDw and ICC measurements obtained from single-echo QSM, multi-echo QSM and R2* maps in the atlas-based cortical ROIs.

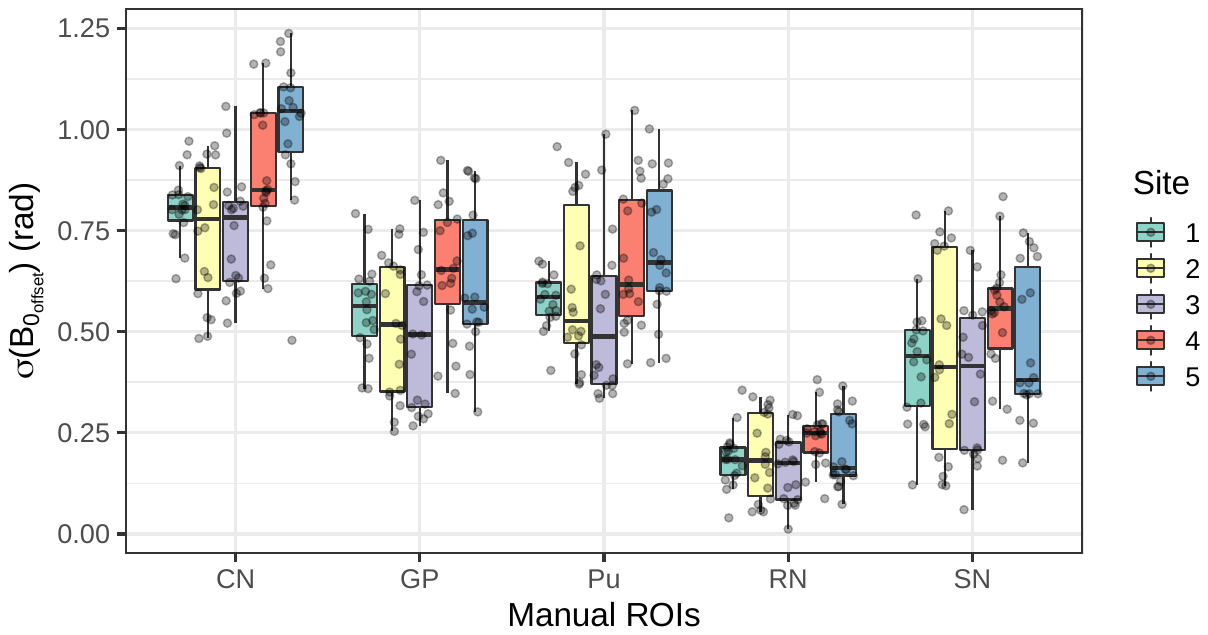

**Figure 3, Supplementary Material 2:** To assess the different shimming conditions across the analysed regions, the standard-deviation of the field-maps (obtained from the background field removal of the multi-echo dataset) were computed within each ROI, $\sigma\left( {B_{0}}_{offset} \right)$. The figure shows boxplots of $\sigma\left( {B_{0}}_{offset} \right)$ in the manual ROIs grouped by sites. Each point in the boxplot represents a subject measurement in the corresponding site. The $\sigma\left( {B_{0}}_{offset} \right)$ was significantly different in the CN (Kruskal-Wallis test, p < 0.001), followed by the Pu and GP ROIs (Kruskal-Wallis test, p=0.036 and p=0.036, respectively), whereas for the other manual ROIs this effect did not reach significance (Kruskal-Wallis test, SN: p=0.15; RN: p=0.11).

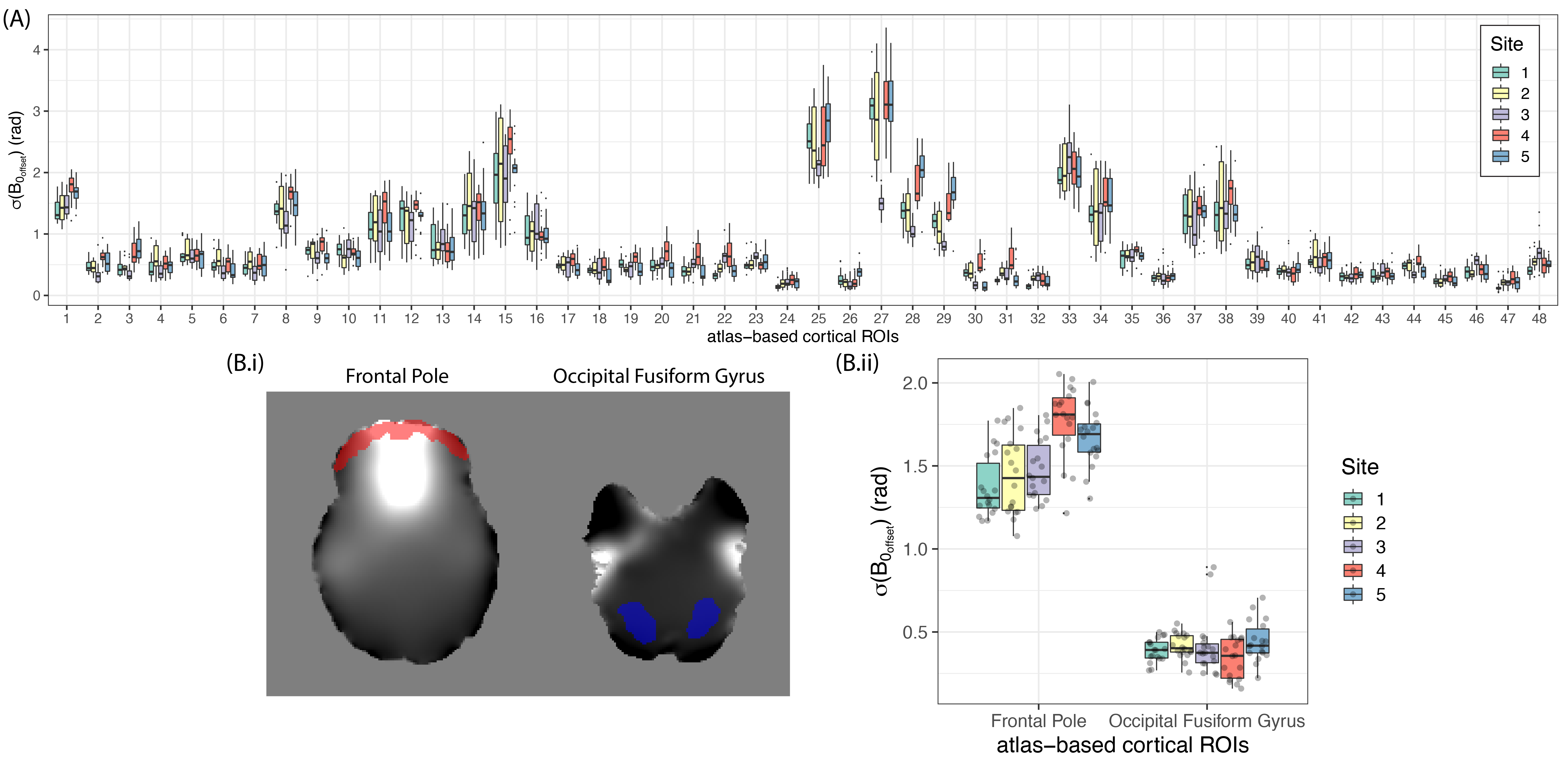

**Figure 4, Supplementary Material 2:** To assess the different shimming conditions across the analysed regions, the standard-deviation of the field-maps (obtained from the background field removal of the multi-echo dataset) were computed within each ROI, $\sigma\left( {B_{0}}_{offset} \right)$. In (A) boxplots of $\sigma\left( {B_{0}}_{offset} \right)$ in the atlas-based cortical ROIs grouped by sites are shown. The names of the numbered ROIs are shown in Table 3, supplementary material 2. Two regions from the cortical ROIs were chosen: axial slices of the frontal pole (red – ROI 1) and occipital fusiform gyrus (blue – ROI 40) ROIs are shown in (B.i). Same plots as in (A) are shown in (B.ii) of the two cortical regions. Each point in the boxplot represents a subject measurement in the corresponding site. In the atlas-based cortical ROIs, regions that showed large variations of the magnetic field (e.g. ROI 1) showed significant cross-site variability (Kruskal-Wallis test, p<0.0001), compared to regions of low B0 field variations (e.g. ROI 40) which did not show significant cross-site variability (Kruskal-Wallis test, p=0.15).

| **number** | **name** |
| --- | --- |
| 1 | Frontal Pole |
| 2 | Insular Cortex |
| 3 | Superior Frontal Gyrus |
| 4 | Middle Frontal Gyrus |
| 5 | Inferior Frontal Gyrus, pars triangularis |
| 6 | Inferior Frontal Gyrus, pars opercularis |
| 7 | Precentral Gyrus |
| 8 | Temporal Pole |
| 9 | Superior Temporal Gyrus, anterior division |
| 10 | Superior Temporal Gyrus, posterior division |
| 11 | Middle Temporal Gyrus, anterior division |
| 12 | Middle Temporal Gyrus, posterior division |
| 13 | Middle Temporal Gyrus, temporooccipital part |
| 14 | Inferior Temporal Gyrus, anterior division |
| 15 | Inferior Temporal Gyrus, posterior division |
| 16 | Inferior Temporal Gyrus, temporooccipital part |
| 17 | Postcentral Gyrus |
| 18 | Superior Parietal Lobule |
| 19 | Supramarginal Gyrus, anterior division |
| 20 | Supramarginal Gyrus, posterior division |
| 21 | Angular Gyrus |
| 22 | Lateral Occipital Cortex, superior division |
| 23 | Lateral Occipital Cortex, inferior division |
| 24 | Intracalcarine Cortex |
| 25 | Frontal Medial Cortex |
| 26 | Juxtapositional Lobule Cortex (formerly Supplementary Motor Cortex) |
| 27 | Subcallosal Cortex |
| 28 | Paracingulate Gyrus |
| 29 | Cingulate Gyrus, anterior division |
| 30 | Cingulate Gyrus, posterior division |
| 31 | Precuneous Cortex |
| 32 | Cuneal Cortex |
| 33 | Frontal Orbital Cortex |
| 34 | Parahippocampal Gyrus, anterior division |
| 35 | Parahippocampal Gyrus, posterior division |
| 36 | Lingual Gyrus |
| 37 | Temporal Fusiform Cortex, anterior division |
| 38 | Temporal Fusiform Cortex, posterior division |
| 39 | Temporal Occipital Fusiform Cortex |
| 40 | Occipital Fusiform Gyrus |
| 41 | Frontal Operculum Cortex |
| 42 | Central Opercular Cortex |
| 43 | Parietal Operculum Cortex |
| 44 | Planum Polare |
| 45 | Heschl's Gyrus (includes H1 and H2) |
| 46 | Planum Temporale |
| 47 | Supracalcarine Cortex |
| 48 | Occipital Pole |

**Table 3, Supplementary Material 2:** Convention for the numbering and given name of the cortical ROIs from the Harvard-Oxford Atlas.

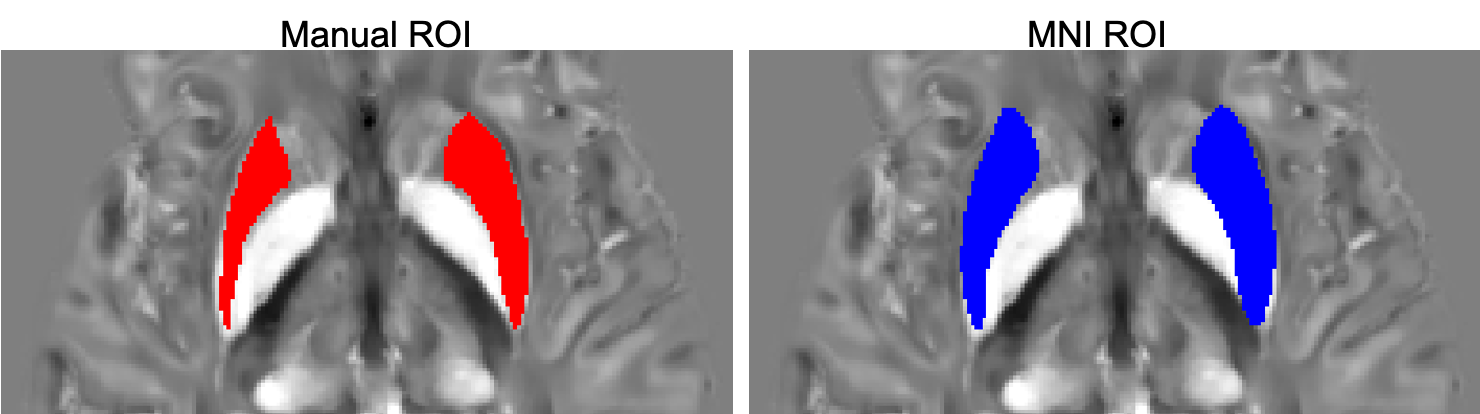

**Figure 5, Supplementary Material 2:** Axial slice showing the putamen ROI overlaid on $\chi$ maps obtained from the manual method (red) and the atlas-based method (blue) on an example subject.

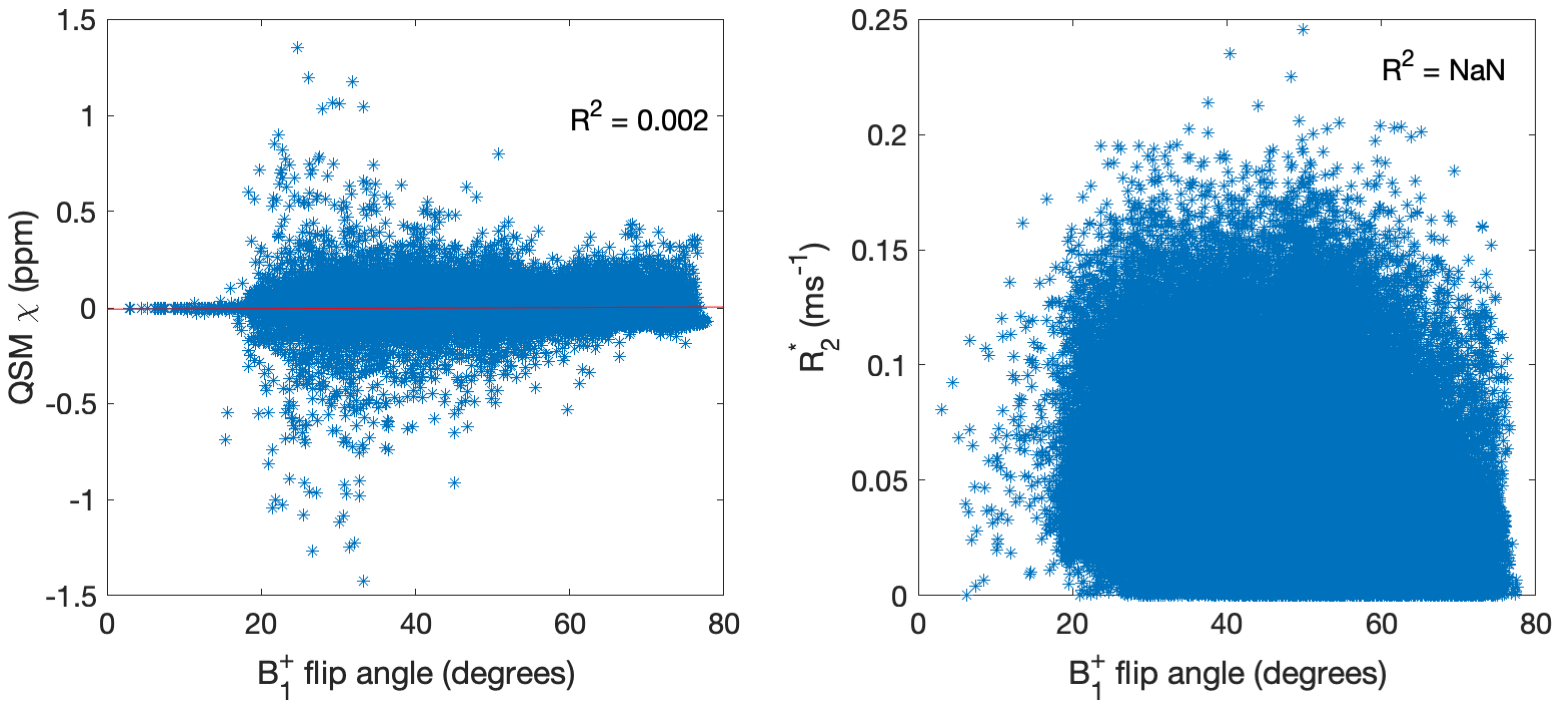

**Figure 6, Supplementary Material 2:** Voxel-wise correlation plots of the transmit B_1_ flip-angle map and the calculated QSM and R2* maps from the multi-echo T2* data from a representative scan.
